# Gene networks associated with early endosperm proliferation and basal endosperm layer differentiation in maize

**DOI:** 10.64898/2025.12.31.697219

**Authors:** Shanshan Zhang, Di Ran, Choong-Hwan Ryu, Guosheng Li, Joanne M Dannenhoffer, Xiangfeng Wang, Gary N. Drews, Ramin Yadegari

## Abstract

Endosperm is a primary source of food and feed globally. Early developmental events including proliferation and cell differentiation in this dedicated seed sink structure are expected be responsive to the availability of photoassimilates and in turn to affect seed size and quality. However, the gene regulatory programs underlying early development of cereal endosperm remain largely unknown. We profiled the maize (*Zea mays*) endosperm transcriptome during the first four days after pollination using laser-capture microdissection and identified multiple temporal co-expression modules including a subset activated upon fertilization. Our analysis of the associated *cis*-regulatory elements and co-expressed transcription factor (TF) families enabled us to construct a gene network regulating basal endosperm transfer layer (BETL) differentiation through MYB-related (MYBR) transcription factors and a network of E2F TFs for early endosperm proliferation. We found a significant association of the BETL network with kernel size variation and a contribution of the Glucose-target-of-rapamycin (TOR)-dependent processes to the E2F-mediated regulation of endosperm proliferation. Using the available data, we propose early proliferative development and transfer cell differentiation in endosperm are coordinated via sugar-sensing inputs to ultimately control seed size. Our analysis can guide the development of strategies for improvement of seed composition and yield.

## INTRODUCTION

How cellular proliferation and early cell fate determination processes are coordinated during early seed development, especially within the nutritive structures of the seed, remain a major unanswered question in plant biology (Liu et al 2022 ARPB; Ref 2). Seed development in angiosperms commences in the female gametophyte with the double fertilization of the haploid egg and the homodiploid central cell to produce a diploid embryo and a triploid endosperm, respectively (Olsen, 2004; Becraft and Gutierrez-Marcos, 2012; Doughty et al., 2014; Lafon-Placette and Kohler, 2014). Early endosperm development is similar in most eudicots and monocots. It is characterized by a series of rapid, coenocytic divisions, followed by a period during which the existing endosperm nuclei cellularize and simultaneously produce additional daughter nuclei which in turn undergo cellularization (Olsen, 2004; Becraft and Gutierrez-Marcos, 2012; Leroux et al., 2014). Subsequently, endosperm development diverges in eudicots and monocots. In most eudicots, the endosperm is nearly absorbed by the developing embryo and cotyledons develop to form the primary seed storage compartment (Olsen, 2004; Lafon-Placette and Kohler, 2014; Liu et al., 2022; Wu et al., 2022). In monocots, the cellularized endosperm undergoes many rounds of mitosis to produce a highly specialized nutritive structure that supports seedling development (Olsen, 2004; Becraft and Gutierrez-Marcos, 2012; Liu et al., 2022; Wu et al., 2022). In maize (*Zea mays*) and other cereals, the central regions of the endosperm undergo endoreduplication, presumably to enhance the ability of the cells to accumulate storage proteins and starch necessary for proper germination (Sabelli and Larkins, 2009). As in other eukaryotes, cellular proliferation and transition to an endocycle in maize endosperm is regulated by a RETINOPBLASTOMA-RELATED (RBR)-E2F network (Sabelli and Larkins, 2009; Desvoyes and Gutierrez, 2020). However, the nature of the transcriptional networks responsible for proliferation of the coenocytic and early cellularized endosperm remain elusive in cereals.

Key cellular differentiation events of the maize endosperm, including formation of the basal endosperm transfer layer (BETL), the aleurone, the starchy endosperm, and the embryo-surrounding region, can be seen as early as 4 days after pollination (DAP) (Olsen, 2004; Becraft and Gutierrez-Marcos, 2012; Leroux et al., 2014; Liu et al., 2022; Wu et al., 2022). The BETL is the primary transfer cell layer of the endosperm and mediates transport of sugar and other metabolites into the endosperm from the underlying maternal kernel tissues (Olsen, 2004; Costa et al., 2012; Liu et al., 2022; Wu et al., 2022). In addition to its role as a sink structure, endosperm has been shown to regulate proper embryogenesis through specific signaling processes during early seed development (Olsen, 2004; Becraft and Gutierrez-Marcos, 2012; Doughty et al., 2014; Lafon-Placette and Kohler, 2014). Endosperm also plays a critical role in regulating seed development, by mediating interactions with the sporophytic maternal cell layers that form the seed coat (Becraft and Gutierrez-Marcos, 2012; Doughty et al., 2014; Lafon-Placette and Kohler, 2014).

By enabling the maternal plant to exert reproductive control over progeny development, the endosperm has likely played a key role in the evolutionary success of the angiosperms (Olsen, 2004; Becraft and Gutierrez-Marcos, 2012). From an economic perspective, the bulk of our agricultural production for food and feed is based on the formation of endosperm in the three main grain crops of maize, rice and wheat (FAO, 2012; Khoury et al., 2014). Consequently, considering its biological and agricultural importance, an understanding of endosperm gene networks including those for proliferation and cell differentiation is important. Although we know many of the players involved in transcriptional regulation of cellular proliferation and endocycle; These networks must integrate developmental and physiological inputs, including the availability of photoassimilates, to regulate endosperm formation and ultimately seed size. As a corollary, whole-plant physiological responses to environmental stresses such as drought likely can affect seed yield by perturbing the same gene networks. Here we report analysis of early maize endosperm gene expression networks through transcriptome profiling of laser-capture microdissected (LCM) endosperm between 0 and 4 DAP. This time series includes the mature haploid central cell (0 DAP), transition to a rapidly proliferating coenocyte upon fertilization (1 and 2 DAP), the coenocyte-to-cellular transition stage (3 DAP), and the fully cellularized endosperm that shows an overall polarity and indications of early cellular differentiation (4 DAP) (Figure 1A and Supplementary Figure 1) (Leroux et al., 2014; Li et al., 2014). Using computational tools, we identified distinct temporal co-expression modules, their co-expressed transcription factor (TF) genes and the associated *cis*-regulatory elements. We found evidence for regulation of a BETL gene network via a set of closely related MYB-related (MYBR) TFs at 4 DAP and found this network to be associated with regulation of seed size. We also detected and experimentally confirmed two E2F-regulated networks with high representation of genes likely regulated by Glucose-target-of-rapamycin (TOR)-dependent (Glc-TOR). Our results support a model in which early-endosperm gene networks associated with BETL development (a cell type responsible for sugar uptake) can influence the size of the developing endosperm and that the proliferation of the endosperm cells via E2F gene networks is regulated via a sugar-sensing pathway.

**Figure 1.**
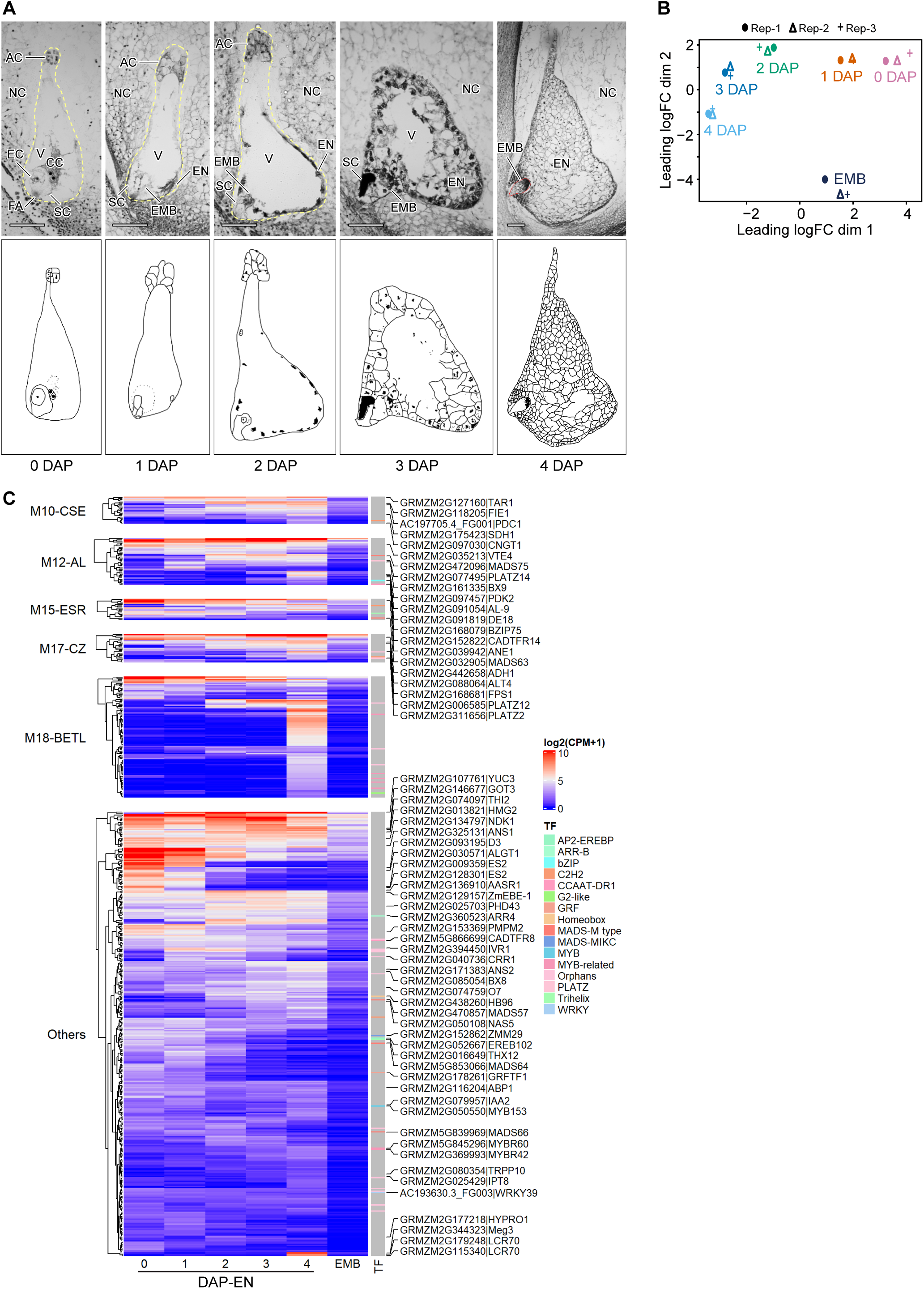
Staging and analysis of mRNA populations of early endosperm in maize inbred B73. (A) Early endosperm time series from kernels at 0, 1, 2, 3, and 4 DAP stained with Toluidine blue O (top) and schematic highlighting of key structural features (bottom). Female gametophytes at 0-2 DAP are outlined in yellow, and whole embryo captured at 4 DAP (EMB) is outlined in red. The micropylar pole is positioned in the lower left portion of each image. AC, antipodal cell; CC, central cell; EC, egg cell; EMB, embryo; EN, endosperm; FA, filiform apparatus; NC, nucellus; SC, synergid cell; V, vacuole. Bars, 100 µm. (B) Visualization of a multidimensional scaling (MDS) of the Euclidean distances of the normalized expression levels for the mRNA populations from an endosperm time series and the embryo for the expressed WGS gene set. Rep-1 to −3, bio-replicate RNAs; 0-4 DAP, endosperm time-series RNAs; EMB, embryo RNA. (C) Relative levels of endosperm preferentially expressed gene mRNAs visualized using a hierarchically clustered heat map based on normalized RNA reads and grouped based on previously identified co-expression modules from 8-DAP kernels (Zhan et al., 2015). Select genes are noted. All associated TF gene families are identified by color-coding.

## RESULTS & DISCUSSION

### Capture, sequencing, and analysis of mRNA populations from early endosperm developmental time series

In order to profile the temporal programs of gene expression in early endosperm development in maize, we isolated and sequenced mRNA populations of inbred B73 endosperm at 0, 1, 2, 3, and 4 DAP, and the whole embryo at 4 DAP (EMB) in three bio-replicates using LCM coupled with RNA-Seq (Figure 1A and Supplemental Figure 1A; Table S1). We aligned ∼768.9 million high-quality reads to the reference genome (Table S2), and further extracted, filtered and normalized the exonic reads (reported as counts per million mapped reads, CPM) based on the annotated gene models (MaizeGDB; Dataset S1). We showed the bio-replicates to be highly correlated (ρ = 0.89 to 0.98), and verified the accuracy of our developmental sampling by interrogating our data using previously described gene expression patterns (Li et al., 2014) (Supplemental Figure 1B; Tables S3). The normalized expression levels not only showed that the individual central cell/endosperm stages cluster distinctly from each other and from EMB, but also showed a clear sequential transition from 0 to 4 DAP (Figure 1B). These changes are likely reflective of temporal changes in mRNA levels as the central undergoes fertilization and early endosperm development.

In order to define endosperm-preferential programs of mRNA accumulation in our samples, we used edgeR (Robinson et al., 2010) to perform generalized linear model (GLM) likelihood ratio tests (3 bio-replicates, FC ≥ 3, FDR ≤ 0.05) for central cell/endosperm stages (collectively termed EN) in comparison to EMB for all 23,867 gene models in FGS (two contrasts in comparison of group means: 0-DAP + 1-DAP + 2-DAP + 3-DAP stage EN vs. 4 × EMB or 4-DAP EN vs. EMB) (Figure 1C; Table S4). Compared to EMB, 558 genes were expressed at significantly higher levels in one or more EN stage(s) (termed EN-pref; Figure 1C; Table S4 ; Dataset S2). Of these, 198 genes were previously reported as belonging to distinct endosperm co-expression modules obtained from an analysis of 8-DAP kernels (Zhan et al., 2015): 22 in central starchy endosperm (M10-CSE module), 38 in aleurone (M12-AL module), 17 in embryo-surrounding region (M15-ESR module), 23 in conducting zone (M17-CZ module), and 98 in basal endosperm transfer layer (M18-BETL module) (Figure 1C; Dataset S2). Using an extensive expression atlas of maize organs including an endosperm stage series (Chen et al., 2014), we found the expression of these genes to display a highly preferential expression pattern within the endosperm (Supplemental Figure 2A). As gene imprinting has been linked with endosperm development (Costa et al., 2012; Gehring, 2013), we evaluated the extent of imprinting programs in the EN-pref gene sets. From a 444-gene-set union of the known imprinted genes previously identified in maize endosperm (Magnard et al., 2003; Gutierrez-Marcos et al., 2004; Waters et al., 2011; Zhang et al., 2011; Xin et al., 2013), we detected 275 imprinted genes as expressed in our dataset (Dataset S3). Among these, we identified 25 maternally expressed genes (MEGs) and 18 paternally expressed genes (PEGs) to be preferentially expressed in the endosperm in our samples (Supplemental Figure 2B). Using the spatial co-expression modules detected in 8-DAP kernels (Zhan et al., 2015), we parsed the imprinted genes in EN-pref and found these genes to display distinct spatiotemporal expression patterns: 4 MEGs (e.g., *Meg1*, *ZMTCRR-1*) and 4 PEGs in BETL, 3 PEGs in CZ, 3 MEGs and 1 PEG in ESR, 3 PEGs (e.g., *MADS75, DE18*) in AL, 6 MEGs (e.g., *FIE1*) and 2 PEGs (e.g., *TAR1*) in CSE (Supplemental Figure 2B). Using edgeR (as described above with GLM likelihood ratio tests, 3 bio-replicates, FC ≤ 1/3, FDR ≤ 0.05), we did a pairwise comparison of individual central cell/endosperm stages vs. EMB (0-DAP vs. EMB, 1-DAP vs. EMB, etc.) to identify EMB-upregulated genes. Intersection of those genes allowed us to identify 275 genes (46 TF genes) with higher mRNA accumulation in EMB as compared to any of the five EN stages (termed EMB-pref, Supplemental Figure 2C; Dataset S4). These included *viviparous-1* (*VP1*) and two other related genes belonging to the ABI3-VP1 family with known roles in embryo development in maize and other plants (Supplemental Figure S2C) (McCarty et al., 1989; Dai et al., 2021). The general embryo-preferential pattern of the EMB-pref gene set was further supported (Supplemental Figure 2D) by an analysis of the individual gene expression patterns using the expression atlas (Chen et al., 2014). Together, our LCM transcriptome data were shown to be of sufficient coverage, resolution, and quality to enable an analysis of gene networks in early endosperm.

### Identification of temporal co-expression modules and the related TF genes and *cis*-motifs associated with BETL differentiation and endosperm proliferation

To identify co-expression modules associated with earliest endosperm developmental events, we extracted temporal programs of gene activity using a StepMiner analysis (Sahoo et al., 2007; Li et al., 2014) of 0- to 4-DAP endosperm. We determined one-step and two-step transitions in RNA levels and identified 20 co-expression modules of four general patterns with varying numbers of genes and TF-gene fraction in each module (*P* ≤ 0.05; Figures 2A, 2B and Supplemental Figures 2 and 3; Dataset S5). Modules were labeled based on which stage(s) displayed an upregulated pattern of RNA levels compared to other stages in the same module (Figure 2B). A majority (80.6%) of the identified genes displayed a single transition point (37.5%, up, 43.1% down) with the rest exhibiting two-step transitions (11.3% up-down, 8.1% down-up; Figure 2A and Supplemental Table 7). The fraction of TF genes varied significantly among the temporal modules with M014, M1, M04 and M4 showing over-representation, whereas M23 and M234 exhibited under-representation of TF genes (*P* < 0.01; Figure 2B). Gene ontology (GO) analysis showed that the StepMiner modules are enriched in unique and shared GO terms (FDR < 0.01; Supplemental Figure 4) that corresponded closely to the expected biological processes. For example, the dominant GO term “cell cycle” in the M1234 module and the GO term “cell-cell signaling” in the M4 module matched with the key developmental events of endosperm proliferation during 1-4 DAP and early endosperm differentiation at 4 DAP, respectively (Supplemental Figure 4) (Leroux et al., 2014).

**Figure 2.**
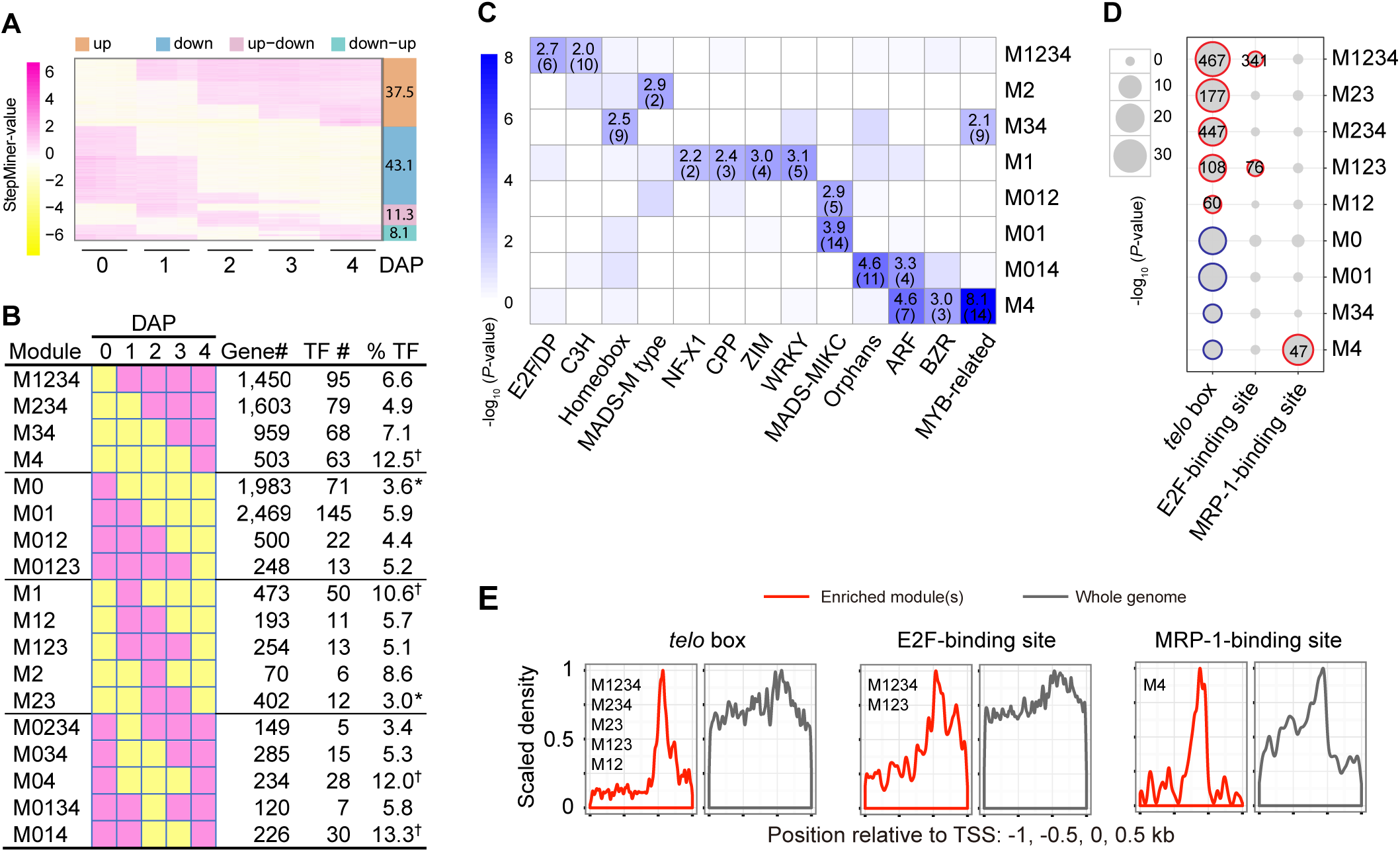
Identification of temporal co-expression modules, and the associated TF genes and *cis*-motifs. (A) Profiles of temporally co-expressed genes of FGS in one-step and two-step transitions for the five stages of endosperm development determined using StepMiner. Yellow denotes low and pink denotes high mRNA levels. Fractions of mRNAs for each of the four general patterns detected are shown as colored bars. (B) A summary table of the characteristics of the main temporal modules including individual patterns, the corresponding gene numbers, the TF gene numbers and fractions thereof. Color coding same as in (A). For ease of notation, module designations only reflect the “up” stages. For example, M123 indicates up-regulation at 1 DAP and down-regulation at 4 DAP. ‘†’ denotes significantly high, ‘*’ denotes significantly low (P < 0.01). The two smallest modules (M3,10 genes; M0124, 5 genes, neither with any TFs) are not shown. (C) Enrichment of the TF gene families in the temporal modules visualized using a hierarchically clustered heat map of –log_10_-transformed *P*-values. The numbers in each box indicate the -log_10_*P*-value and the numbers of genes (in parenthesis) for each TF family (*P* < 0.01). Analysis of all TF gene families (59) in all temporal modules is shown in Dataset S6. *P*-values were calculated using Fisher’s exact test. (D) Enrichment of the *cis*-motifs in the temporal modules visualized using a scatter plot of –log_10_-transformed *P*-values. Over-representation and under-representation (*P* < 0.001) are circled by red and blue lines, respectively. The numbers in each cell indicate the numbers of genes containing a *cis*-motif with a cutoff of *P* < 0.001 (over-representation). *P*-values were calculated using two-sided Fisher’s exact test. (E) Scaled density plots of the three enriched *cis*-motifs across −1 kb to +0.5 kb (relative to annotated TSS) of genes in the enriched module(s) or whole genome.

**Figure 3.**
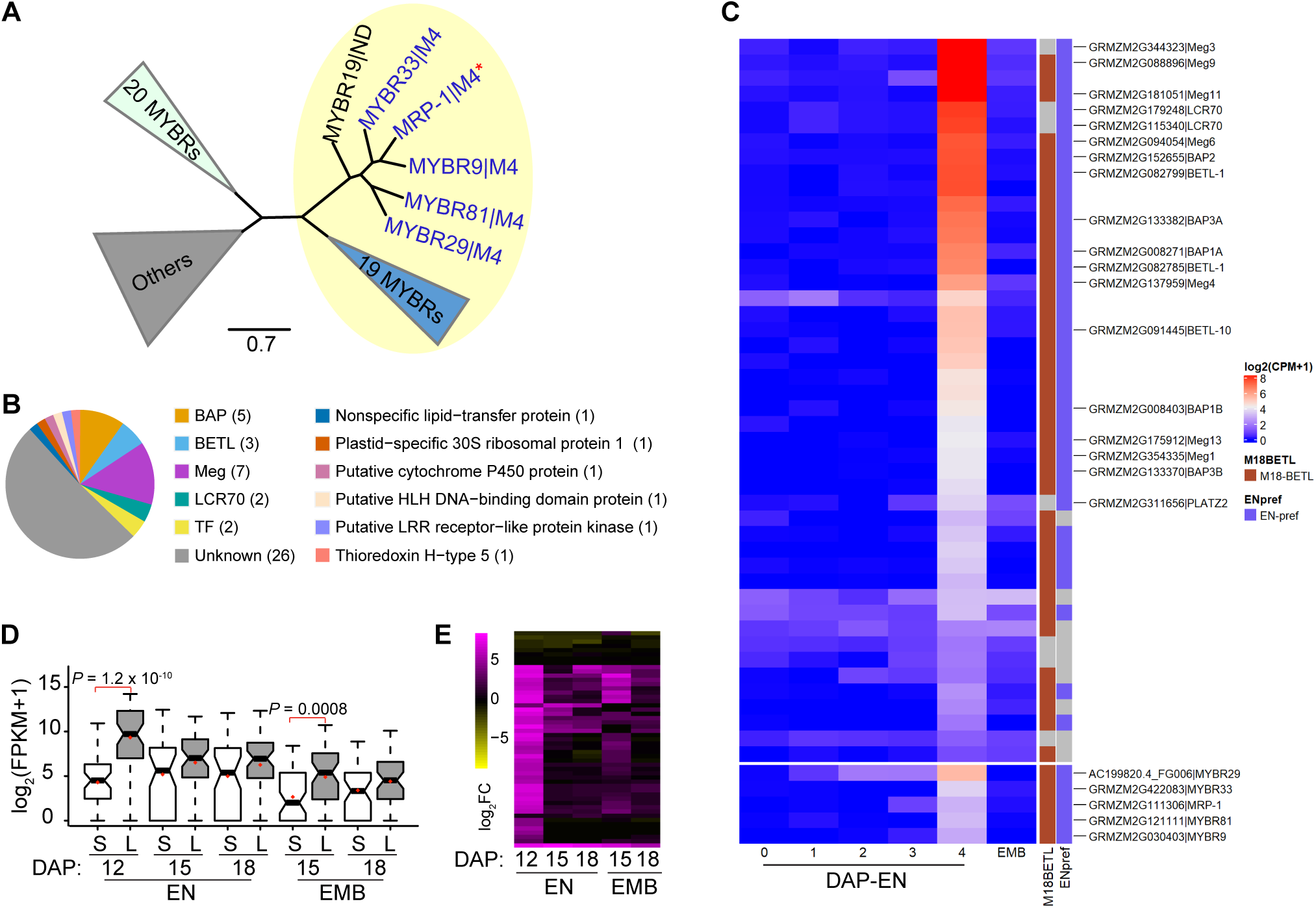
Identification of a BETL-associated regulatory module in early maize endosperm. (A) Phylogenetic analysis of MYBR TF family in maize using predicted amino acid sequences. Five members of the MRP-1 subclade encoding MRP-1, and MYBR9, 29, 33, and 81 were detected as expressed in the M4 module (blue font/asterisk). All 25 proteins in the clade highlighted in yellow and 13 of 20 proteins in the clade highlighted in green have a highly conserved SHAQK(Y/F)F amino acid domain. The maximum likelihood tree was constructed using the program RAxML(Stamatakis et al., 2008). ND denotes not detected as expressed in our dataset. Asterisk (*) indicates the previously described MYBR TF gene MRP-1. Bar, 0.7 amino acid substitutions per site. A detailed tree is provided in Supplemental Figure 5. (B) Gene annotations for M4-47 genes based on the available data and using EnsemblPlants. Numbers of individual genes encoding each protein type is indicted in parentheses. (C) The developmental mRNA abundance levels of M4-47 and the MYBR TF genes shown using a hierarchically clustered heat map. Genes previously identified as co-expressed in a BETL module (M18-BETL Zhan et al 2015 and/or as preferentially expressed in endosperm (EN-Pref) in this study are color coded in separate columns. Select genes including all five expressed MYBR TF genes are identified. (D) Differential distribution of the normalized RNA reads of the M4-47 genes using the reported expression atlas of the KLS30 (L) vs. KSS30 (S) kernels (Sekhon et al., 2014) shown as boxplots. Bold bars indicate median, and red diamonds indicate mean values. Only *P*-values with a cutoff of *P* < 0.01 are indicated. *P*-values were obtained using a Wilcoxon rank-sum test. (E) Hierarchically clustering analysis of mRNA levels the M4-47 genes based on log_2_-transformed fold-change (FC) of the KLS30 vs. KSS30 kernels (Sekhon et al., 2014). EN, endosperm; EMB, embryo.

To identify the major co-expressed TFs that may function in these temporal modules, we tested each module for over-representation of annotated maize TFs (Yilmaz et al., 2009; Jin et al., 2014) and showed that 13 of the 59 maize TF gene families were significantly enriched in at least one module (*P* < 0.01; Figure 2C; Dataset S6). The ARF genes were over-represented in M014 (4 genes) and M4 (7; Figure 2C) suggesting an active role for auxin-regulated programs during endosperm proliferation and differentiation, and in agreement with previous observations in maize and Arabidopsis (Chen et al., 2014; Locascio et al., 2014; Köhler et al., 2021). In addition, the MADS-MIKC family genes were found to be over-represented in M01 (14) and M012 (5), whereas two MADS-M type genes were highly enriched in M2 (Figure 2C). These distinct association patterns imply that the MADS-MIKC TFs function preferentially before or during fertilization while the MADS-M TFs may function during coenocytic endosperm development as has been described previously for Arabidopsis and rice (Gramzow and Theissen, 2010; Masiero et al., 2011). In contrast to other modules, M1 displayed both a high proportion of TF genes (10.6%) and a broad representation of TF gene families including NF-X1, WRKY, CPP and ZIM (Figure 2B and 2C). The latter two families have previously been shown to function in reproductive development and cell division (Liu et al., 1997; Hauser et al., 1998; Hauser et al., 2000; Song et al., 2000; Sijacic et al., 2011), and regulation of jasmonic acid signaling during fertilization in plants (Wasternack et al., 2013), respectively. Therefore, the M1 module likely marks a key transition from the gametophytic to the zygotic state upon fertilization. Relevant to our study here, we detected 15 MYB-related (MYBR) TF genes including *MRP-1*(*Myb-Related Protein-1*) as being over-represented in M4, and detected a significant enrichment for the E2F/Dimerization Partner (DP) genes in M1234 (Figure 2C). These two TF families can be linked with corresponding *cis*-motifs that were overrepresented in their respective co-expressed gene sets (Figure 2D, see below). MRP-1 has been linked with BETL cellular differentiation in maize via activating a hypothesized gene network through its cognate *cis*-motifs (MRP-1 binding sites flanking its target genes) (Gomez et al., 2002; Barrero et al., 2006; Gomez et al., 2009; Zhan et al., 2015). E2F family members play a conserved key role in transcriptional regulation of genes required for G1-S transition in animals and plants (Berckmans and De Veylder, 2009; Bertoli et al., 2013; Dante et al., 2014; Gutierrez, 2016; Sablowski and Gutierrez, 2022). In Arabidopsis, overexpression of E2Fa and DPa can drive differentiated non-dividing cells to re-enter S-phase in cell cultures and *in planta* (De Veylder et al., 2002; Rossignol et al., 2002), indicating that the transcriptional upregulation of *E2F* genes can be associated with G1-S transition in plants. Together, these preliminary data suggest that we have detected distinct co-expressed gene sets associated with BETL cell differentiation and endosperm proliferation. Therefore, we focused on further analysis of the transcriptional networks regulated by MRP-1/MYBR and E2F TFs in this study.

To detect the putative target gene sets related to the co-expressed TFs, we first identified 10-12-bp sequence motifs over-represented within −1 kb to +0.5 kb (relative to the annotated transcription start site, TSS) of the genes in each module using MEME (Bailey et al., 2006) (Dataset S7) and further narrowed down the putative DNA-binding *cis*-regulatory motifs (*cis*-motifs) by finding those most significantly similar to the previously described plant *cis*-motifs (Mathelier et al., 2014) using the Tomtom program (Gupta et al., 2007) (Dataset S7). This analysis uncovered MRP-1-binding sites in M4 and *telo* (short interstitial telomere) boxes enriched in modules M1234, M234, M12, M123 and M23 (Dataset S7). The former is consistent with the detection of *MRP-1* in M4 (Figure 2C) and supports the hypothesis that an MRP-1-regulated network is active as early as 4 DAP (see below). On the other hand, rather than detecting E2F-binding sites, our MEME analysis uncovered *telo* boxes, likely due to their higher prevalence and sequence conservation as compared to the known E2F-binding sites. *telo* boxes have been shown to mediate transcriptional activation of genes encoding components of the translational machinery in rapidly dividing cells in Arabidopsis and rice (Tremousaygue et al., 1999; Tremousaygue et al., 2003; Gaspin et al., 2010), often in co-occurrence with E2F-binding sites (Tremousaygue et al., 1999; Rossignol et al., 2002; Tremousaygue et al., 2003; Gaspin et al., 2010), and as *cis*-motifs for recruitment of Polycomb repressive complexes in Arabidopsis (Zhou, 2016; Xiao, 2017; Zhou, 2018; Bieluszewski, 2021). Consequently, we used PatMatch (Yan et al., 2005) to search for occurrence of the MEME-detected motifs upstream of all genes in the genome and determined their enrichment in each temporal module (*P* < 0.001; Figure 2D; Table S5; Dataset S8). As expected, MRP-1 binding sites showed significant enrichment in M4 but not in other modules (Figure 2D). On the other hand, *telo* boxes were enriched in five modules: M1234 (467 of 1450 genes), M123 (108 of 254 genes), M234 (447 of 1603 genes), M12 (60 of 193 genes), and M23 (177 of 402 genes) (Figure 2D). Modules M1234 and M123 also showed significant enrichment of *E2F* binding sites with 341 and 76 genes containing known E2F-binding sites, respectively (denoted as M1234-341 and M123-76), and 120 and 36 genes also displaying co-occurrence of both *cis*-motifs, respectively (Figure 2D; Dataset S8). These *telo*- or E2F-*telo*-associated modules all shared an up or up-down pattern and were inclusive of stage 2 DAP but were exclusive of 0 DAP (Figure 2D). Likewise, the four modules (M0, M01, M34 and M4) showing significant under enrichment of *telo* boxes were exclusive of stage 2 DAP but inclusive of 0 or 4 DAP (Figure 2D). Both motifs showed a high-density distribution pattern centered near the annotated TSS sequences in the enriched modules (Figure 2E), suggesting roles in transcriptional regulation of their respective downstream genes. Therefore, one or more E2F-*telo*-related regulatory programs likely mediate activation of a large number of genes after fertilization with peaks of expression at 2 DAP. Taken together, our analysis identified early temporal modules associated with BETL cell differentiation and post-fertilization proliferative development of the endosperm.

### An MYBR-regulated network associated with BETL cell differentiation

Because MRP-1 has been linked with BETL cell differentiation, the observed co-enrichment of *MRP-1* (and the related MYBR genes) and genes with MRP-1-binding sites in M4 (Figures 2C and 2D) prompted us to further examine this module for the associated TFs and *cis*-motifs (Supplemental Figure 2). Five of the 15 MYBR genes, including *MRP-1*, and *MYBR9*, *29*, *33*, and *81* formed a tight phylogenetic clade within the larger MYBR gene clade (Figure 3A and Supplemental Figure 5). All five also exhibited the highly conserved SHAQK(Y/F)F amino acid motif within the C-terminus portion of the Myb domain previously reported for MRP-1 (Gomez et al., 2002) (Supplemental Figure 6A). Our *de novo* detection of *cis*-motifs in the M4 module identified all four MRP-1-binding sub-motifs, previously shown to bind MRP-1 (Zhan et al., 2015) (Dataset S7, S9). Using a genome-wide PatMatch (Yan et al., 2005) search, these motifs were found to be preferentially located immediately upstream of the annotated gene models in M4 as compared to their genome-wide distribution (Figure 2E; Dataset S8, S9). We found 47 genes with the MRP-1-binding sub-motifs to be over-represented in the M4 module (*P* < 0.001; denoted as M4-47 genes; Figure 2D; Dataset S9). Among these genes, 43 were previously reported as BETL-specific (Zhan et al., 2015), including 14 encoding BAP (5 genes), BETL (3 genes), and Meg (6 genes) (Figure 3C; Dataset S9). Using the expression atlas (Chen et al., 2014), we detected highly preferential and coordinated expression patterns in the endosperm for both the M4-47 and the five SHAQK(Y/F)F-motif MYBR genes (Supplemental Figure 6B). Both gene sets generally displayed an upregulation pattern starting at 4 DAP and continuous expression in a subsequent stage (Figure 3C and Supplemental Figure 6B). Therefore, the M4-47 genes may contribute to a gene network that acts as early as 4 DAP to drive BETL cell differentiation.

Among the M4-47 genes, the maternally expressed gene *Meg1* was shown to play a role in control of seed size (Costa et al., 2012). To understand the extent of contribution of the M4-47 gene network to the overall development of endosperm and its potential role in controlling seed size, we analyzed a published dataset of endosperm and embryo RNA profiles obtained from two divergently selected maize populations of large (KLS30) and small (KSS30) kernels (Sekhon et al., 2014) and looked for any significant correlation between up- or down-regulated patterns in the endosperm of KLS30 vs. KSS30 seeds with our gene sets (|log_2_fold-change| > 1 at any stage; denoted as KLS30-up or KSS30-down, and KLS30-down or KSS30-up, and abbreviated henceforth as KLS30-up and KLS30-down, respectively). We performed a Wilcoxon rank-sum test of the distribution of the normalized RNA reads for all temporal modules as well as M4-47 genes in the published dataset (12, 15 and 18 DAP for endosperm, and 15 and 18 DAP for embryo) (Sekhon et al., 2014) (Supplemental Figure 7 and Figure 3D). We found the expression of the individual modules M4 as well as M01 and M1234 to show increased expression in KLS30 (vs. KSS30) endosperm at 12 (M01, M4), 15 (M1234) and 18 (M1234) DAP (*P* < 0.001; Supplemental Figure 7). For M4 genes, an increased expression in KLS30 endosperm at 12 DAP (*P =* 8.4 × 10^−7^) could be attributed to the presence of the MRP-1-binding sites in the M4-47 subset (*P =* 1.2 × 10^−10^; Figure 3D and Supplemental Figure 7). Further, we identified the individual genes with the upregulated patterns in the endosperm of KLS30 vs. KSS30 seeds (|log_2_fold-change| > 1 at any stage; Figure 3E) and found 2 KLS30-down genes and 42 KLS30-up genes, including *Meg1* and five *BAPs* and three *BETLs* among the M4-47 genes. This is consistent with the observed phenotype of *Meg1* knock-downs which produce smaller seeds with limited BETL differentiation and reduced sugar content (Costa et al., 2012). While a *CaMV35*S enhancer-driven overexpression of *Meg1* produced small seeds, overexpression studies using BETL-specific promoters produced large-sized seeds in a dosage-dependent manner (Costa et al., 2012). These observations suggested a requirement for proper spatial and quantitative expression of *Meg1* in the BETL to effect proper seed size (Costa et al., 2012; Haig, 2013). Dysregulation of some of the M4-47 genes, including *MRP-1* and *BETL-1*, and an associated defect in BETL have been reported in mutants or genetic lines with excess maternal or paternal genomes (Cooper, 1951; Lin, 1984; Leblanc et al., 2002; Pennington et al., 2008). A genome-wide scan of the maize Krug Yellow Dent and its derivatives, KLS30 and KSS30, identified 94 candidate divergent regions (at 99.9% outlier level) likely under artificial selection for seed size (Odhiambo and Compton, 1987; Russell, 2006; Hirsch et al., 2014). When mapped to the B73 RefGen_v3 (5b) genome annotation in EnsemblPlant, we detected 2417 genes in the selected regions. Of these, 10 were M4-47 genes, including *BAP2*, seven *Meg* genes (including *Meg1*), and 2 unknown genes, with all showing a KLS30-up expression pattern (Dataset S9). Taken together, our data suggest that the M4-47 genes are regulated through MRP-1/MYBR TFs, and in turn impact seed size through regulation of BETL cell differentiation in maize.

### E2F regulation of genes associated with endosperm proliferation

As described above, the M1234 module was shown to be co-enriched with the E2F/DP TFs and the associated E2F binding sites (Figures 2C and 2D). Eighteen of the 19 E2F/DP-coding TF genes annotated in the maize genome were shown to be endosperm-expressed, and six of these genes were found to be co-expressed in the M1234 module (E2F5, E2F6, E2F7, E2F13, E2F18, and E2F19; Figure 4A; Dataset S10). Phylogenetic analysis indicated that these six genes could be grouped into one of the three main canonical E2F subfamilies in maize, including the typical E2Fs (E2F5, most closely related phylogenetically to Arabidopsis E2FC), the DPs (E2F6, E2F18 and E2F19), and the atypical E2Fs (E2F7 and E2F13) (Figure 4B; Dataset S10). Because the E2Fa/b-clade proteins of the typical-E2F subfamily (Figure 4B; Dataset S10) have been reported to interact with DP proteins to regulate gene expression in Arabidopsis and rice (Berckmans and De Veylder, 2009; Bertoli et al., 2013; Gutierrez, 2016), we also included in our following analyses three E2Fa/b clade proteins (E2F4, E2F11 and E2F16) which were co-expressed with DP proteins in endosperm but not in the M1234 module (Figure 4A; Dataset S10). Using Y2H, we subsequently found that the expressed DPs E2F6, E2F18 and E2F19 to be able to interact with the typical E2F proteins E2F4, E2F5, E2F11 and E2F16, but not with the expressed atypical E2F proteins E2F7 and E2F13 (Figure 4C and Supplemental Figure 8).

**Figure 4.**
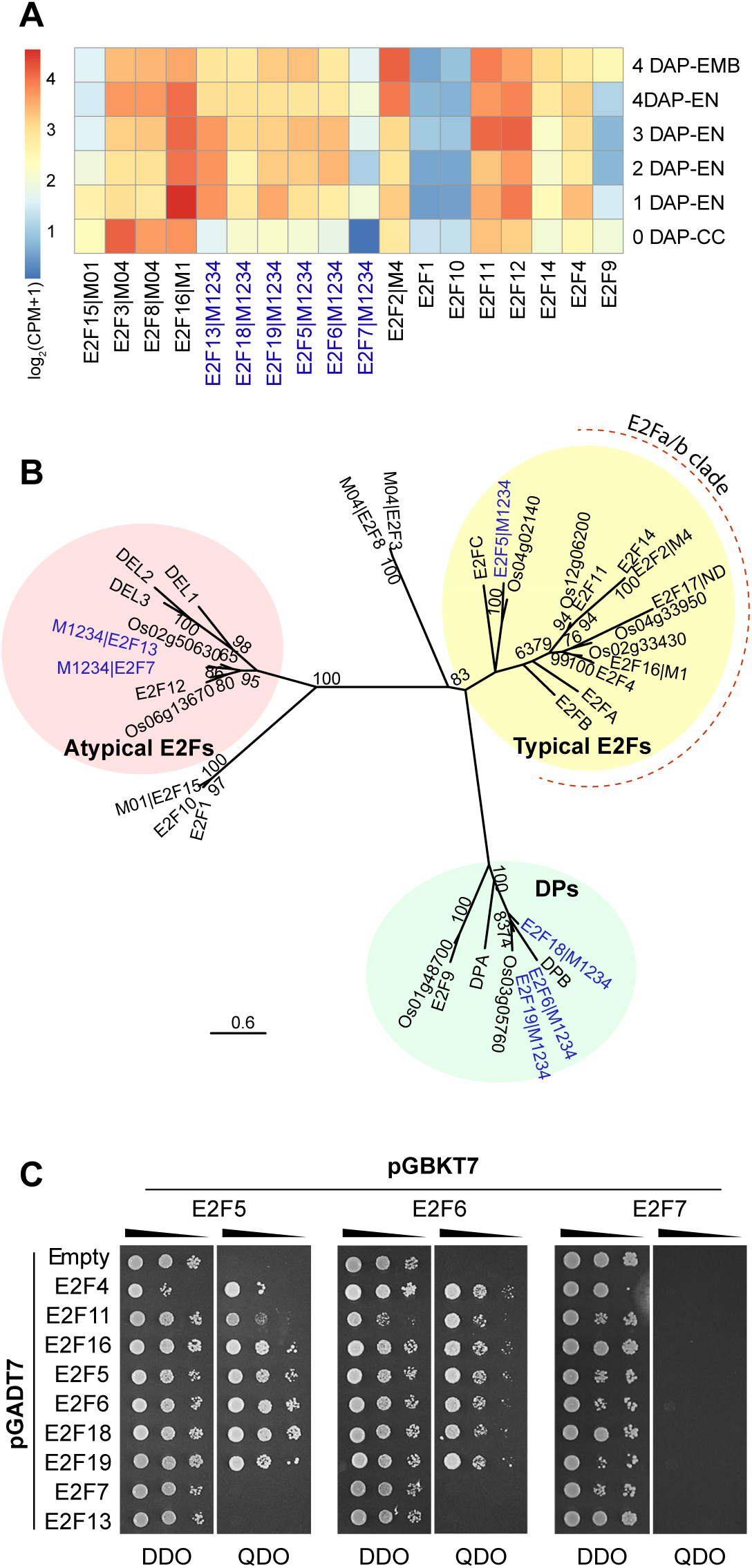
Characterization of the candidate *E2F* TF genes in early endosperm proliferation. (A) The expression patterns of genes encoding E2F proteins using our LCM-RNA-Seq time series. The M1234 module genes are indicated in blue font. (B) Phylogenetic tree of E2F/Dimerization Partner (DP) TF family in maize, Arabidopsis and rice using predicted amino acid sequences. Gene identities were obtained from the available maize (E2F1-19), Rice (LOC_Osxxgxxxxx, “LOC_” omitted), and Arabidopsis (E2FA-C, DEL1-3, DPA and DPB) databases. Where applicable, the temporal-module expression pattern associated with each maize gene is indicated as an extension of the gene name. ND denotes not detected as expressed in our dataset. The genes closely clustered as typical E2Fs, DPs and atypical E2Fs are highlighted in yellow, green, and pink, respectively. The M1234 module genes are indicated in blue font. The maximum likelihood tree was constructed using the program RAxML (Stamatakis et al., 2008). Values above the nodes are from 100 bootstrap replicates; only values >60% are reported. Bar, 0.6 amino acid substitutions per site. (C) Detection of the protein-protein interactions between the candidate E2Fs using yeast two-hybrid analysis. Yeast *Saccharomyces cerevisiae* AH109 cells were co-transformed with the indicated bait and prey combinations. The proteins listed were fused with a Gal4 DNA-binding domain (BD, pGBKT7) or a Gal4-activation domain (AD, pGADT7). Double drop-out (DDO, left), synthetic complete medium lacking Trp and Leu; quadruple dropout (QDO, right), synthetic complete medium lacking Trp, Leu, His, and adenine. Co-transformed colonies growing on QDO were considered as positive interaction transformants. Additional analyses of the protein-protein interactions between the candidate E2Fsare shown in Supplemental Figure 8.

Guided by these protein-protein interaction results, we tested the ability of a representative subset of the endosperm-expressed E2Fs (E2F5, E2F6, E2F7 and E2F16) to bind either as heterodimers or homodimers to a set of detected E2F-binding sites in a dual-luciferase transactivation assay. Among these sites, sub-motif a was the dominant E2F-binding site variant/sub-motif in the M1234 and M123 modules (Supplemental Figure 9A; Dataset S11). The same core sequence upstream of the gene encoding retinoblastoma-related3 (RBR3) protein had been shown to bind an E2F protein *in vitro* (sites I and IV) (Sabelli et al., 2005). Our binding assays showed that E2F5 or E2F16 in combination with E2F6 could bind to the E2F-binding sub-motif a concatemers (3×, 6×) and activate reporter LUC expression (Figure 5A and Supplemental Figure 9B), indicating that E2F5-E2F6 and E2F16-E2F6 heterodimers possess an ability to activate transcription *in vivo*. On the other hand, no significant transactivation was detected in assays with the atypical E2F7 (Figures 5A and Supplemental Figure 9B). Similar results were found with the other variants of the E2F-binding site (Supplemental Figure 9C). Binding of E2F5-E2F6 and E2F16-E2F6 heterodimers to E2F-binding site was shown to be specific as the mutated E2F-binding sub-motif a (6×) could not confer reporter gene transactivation (Supplemental Figure 9D). A series of electrophoretic mobility shift assays (EMSA) further supported the specific binding of E2F16-E2F6 to E2F-binding sites in the promoters of the *PCNA2*, *E2F5*, and *E2F13* genes in M1234 (Supplemental Figure 9E). These results identify the E2F-binding site sub-motifs associated with the M1234 module as potential targets of the E2F proteins in early endosperm.

**Figure 5.**
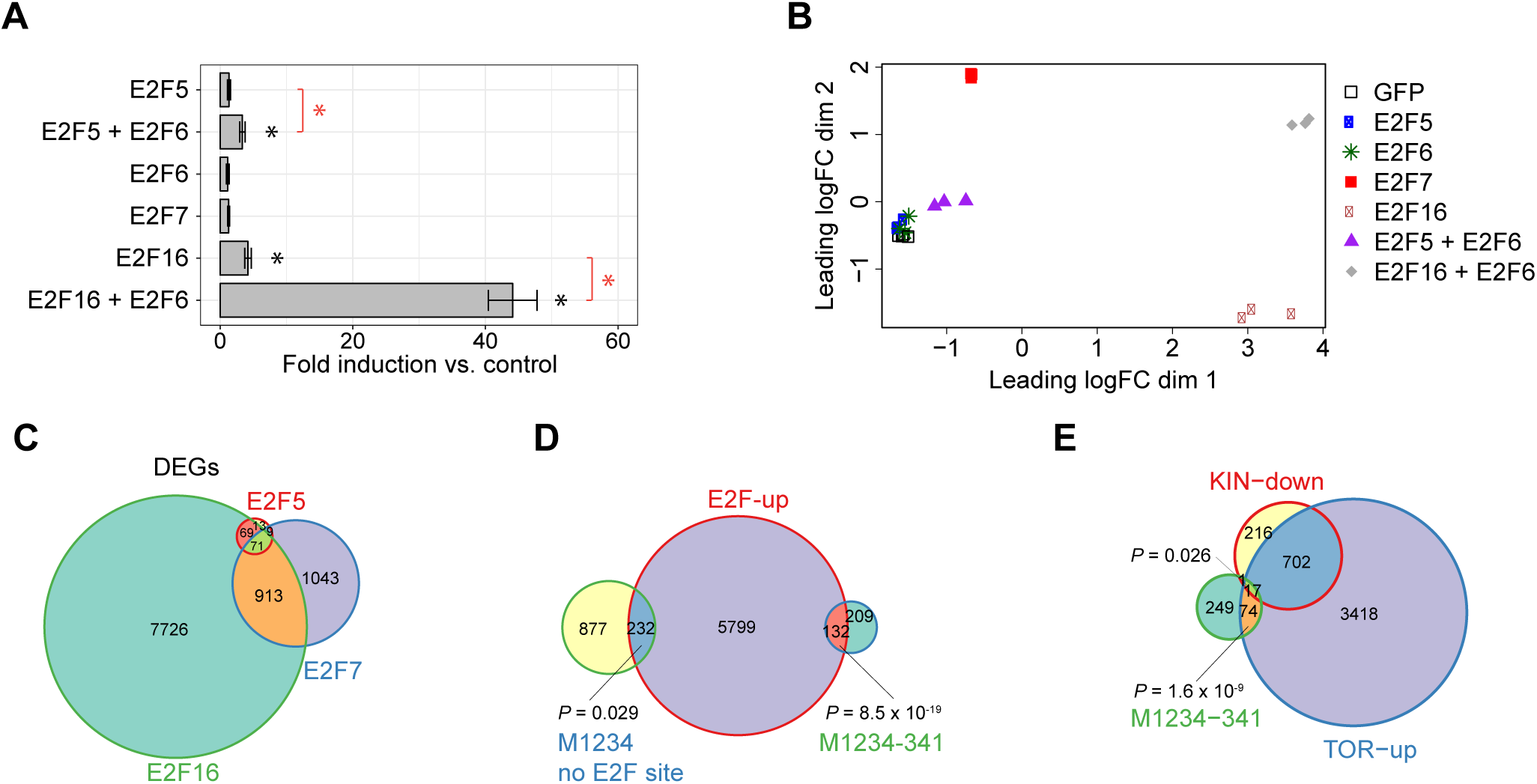
Early-endosperm regulatory E2Fs and activation of their target genes. (A) Dual-luciferase transactivation assays for detection of single E2F and combination of E2Fs binding to a concatemerized (6×) E2F-binding *cis*-motif (submotif a, see Supplemental Figure 9A for details). Detected binding is measured as fold induction vs. control (mean ± SEM; *n* = 5; see Methods for details). Asterisks indicate *P* < 0.05, Student’s *t* test. (B) Multidimensional scaling (MDS) of the Euclidean distances of the normalized expression levels for the mRNA populations from RNA-Seq of maize mesophyll protoplasts with transient expression of single or combination of *E2F*s (3 replicates each) or *GFP* (control, 4 replicates). (C) Intersection of DEGs (vs. control GFP, FDR < 0.05 with |log_2_FC| > log_2_(1.5)) due to transient ectopic expression of E2F5, E2F7, and E2F16 in maize mesophyll protoplasts shown using a Venn diagram. (D) Intersection of all ectopic E2F-upregulated genes from above (E2F-up, see Supplemental Figure 10A), M1234-341, and M1234 genes without E2F-binding sites in the upstream flanking regions, within −1 kb to +0.5 kb relative to annotated TSS, shown using a Venn diagram. (E) Intersection of M1234-341 genes, TOR-up genes, and KIN10-down genes shown using a Venn diagram. *P*-values were calculated using Fisher’s exact test. Detailed information on the TOR-up/down genes is provided in Supplemental figure 11.

To further understand the extent of the gene networks regulated by these E2F proteins, we transiently expressed E2F5, E2F6, E2F7, and E2F16 individually, and in combinations of E2F5 + E2F6, and E2F16 + E2F6 in maize mesophyll protoplasts and profiled the transcriptomes. We detected distinct changes in the mRNA populations as a result of ectopic expression of the E2Fs (Figure 5B; Dataset S12). We used edgeR to identify differentially expressed genes (DEGs, E2F(s) vs. control GFP; FDR < 0.05 with |log_2_FC| > log_2_(1.5); Figure 5C and Supplemental Figure 10A; Dataset S13) and found that the dimerization partner E2F6 can synergistically activate target genes with either E2F5 or E2F16 (Supplemental Figures 10B and 10C). This agrees with the results of our transactivation and DNA-binding assays with select target genes (Figure 5A and Supplemental Figure 9) and is in line with previous observations where DPs in combination with typical E2Fs can synergistically activate target gene expression (Kosugi and Ohashi, 2002; Rossignol et al., 2002). In fact, the union set of E2F-upregulated DEGs (6163 E2F-up genes; Supplemental Figures 10A; Dataset S13) constituted more than a third of the presumptive E2F target gene subsets of the temporal modules of interest M1234 (M1234-341, 132/341, 38.7%) and M123 (M123-76, 32/76, 42.1%) (Figure 5D and Supplemental Figure ; Dataset S11). In addition, we detected another 232 genes from M1234 and 46 genes from M123 (20.9% and 25.8% of all genes, respectively) containing no detectable E2F-binding sites to be upregulated by E2Fs in the protoplasts (Figure 5D and Supplemental Figure 10D; Dataset S13), likely corresponding to indirect targets of E2Fs. Among E2F genes, we found that the E2F16 + E2F6 combination can upregulate expression of E2F5, E2F7 and E2F13 in maize mesophyll protoplasts (Supplemental Figure 10E), indicating a complex multiple-layer regulation with the feedback transcriptional activation of E2F genes themselves. Taken together, our data identifies a network of E2F-regulated genes in post-fertilization proliferative development of maize endosperm.

### Association of early endosperm gene networks with glucose-TOR pathway

In response to nutrient status, E2F proteins have been reported to be phosphorylated by a glucose-TOR signaling pathway to regulate target gene expression and further regulate the G1-S transition and promote cell cycle initiation in Arabidopsis meristems (Xiong et al., 2013; Liu and Xiong, 2022). We detected the expression of several genes encoding invertases, sucrose and hexose transporters, and the TOR complex (TORC) proteins annotated in MaizeGDB (Lawrence et al., 2004) in early endosperm (Supplemental Figure 11A; Dataset S14). Accordingly, we hypothesized that a subset of the gene expression programs identified here would be responsive to TOR kinase-signaling processes (Laplante and Sabatini, 2012; Sheen, 2014), which would sense and integrate nutrient and other cues to regulate early endosperm proliferation. To address this, we tested all temporal modules as well as the M1234-341 and M123-76 subsets containing E2F-binding sites (Dataset S11) for expression of genes previously identified in Arabidopsis seedlings to be up- or down-regulated by a physiological concentration of glucose (Glc) through a TOR signaling pathway (Xiong et al., 2013) (denoted as Glc-TOR-up and Glc-TOR-down genes, respectively; Supplemental Figure 11B; Dataset S14). We found a high representation of Glc-TOR-up genes in modules M1234, M234, M12, M123, M23 and M0123, and conversely, a high representation of Glc-TOR-down genes in three modules all with an upregulated 0-DAP pattern (M0, M01, M014; Supplemental Figures 11C; *P* < 0.001). Furthermore, the former Glc-TOR-up genes were generally associated with cellular functions such as protein synthesis, cell cycle and DNA synthesis whereas the Glc-TOR-down genes were associated with protein degradation and autophagy (Laplante and Sabatini, 2012; Xiong et al., 2013; Sheen, 2014) (Supplemental Figure 11D; Dataset S15; *P* < 0.001). The detected Glc-TOR-up/down gene sets in Arabidopsis were nearly fully inclusive of genes shown to be repressed or activated by ectopic expression of the *PROTEIN KINASE10* (*KIN10*, the catalytic subunit of energy sensor kinase) in Arabidopsis protoplasts (Baena-Gonzalez et al., 2007; Sheen, 2014) (denoted as KIN10-down and KIN10-up, respectively; Supplemental Figure 11E). Because ectopic expression of *KIN10* induces expression of several autophagy genes while simultaneously suppresses genes involved in protein synthesis and anabolism (Baena-Gonzalez et al., 2007), Glc-TOR-up genes are expected to correlate with KIN10-down genes, and Glc-TOR-down genes correlate with KIN10-up genes (Sheen, 2014). Accordingly, we looked for homologs of the same genes in our datasets and found a high representation of Glc-TOR-up genes (but not Glc-TOR-down genes) and KIN10-down genes in M1234-341 (91/341 with *P* = 1.6 × 10^−9^ and 18/341 with *P* = 0.026, respectively; Figure 5E), and M123-76 (27/76 with *P* = 2.8 × 10^−6^ and 10/76 with *P* = 1.1 × 10^−4^, respectively; Supplemental Figure 11F). This supports the hypothesis that early endosperm proliferation in maize involves glucose-sensing, TOR-dependent transcriptional responses. Together, our results suggest that upon fertilization a major reprogramming of the central cell, through a series of Glc-TOR-activated (and KIN10-suppressed) processes, underlies rapid endosperm proliferation through the E2F-regulated networks.

## CONCLUSIONS

In this study, we provide a comprehensive analysis of the maize endosperm transcriptome from the earliest post-fertilization stages of kernel development. Using temporal co-expression modules and the related TF genes and *cis*-motifs, we constructed a preliminary set of gene networks associated with the two key biological processes of early endosperm proliferation and cell differentiation. Endosperm proliferation occurs initially in response to fertilization and both the extent of coenocytic and cellular proliferation processes are thought to be major determining factors of the ultimate endosperm cell number which in turn establishes the sink capacity of the monocot seed (Jones et al., 1996). These processes are expected to be responsive to nutrient status, particularly the availability of maternally supplied sugars within the developing seed (Cazetta et al., 1999; Cheng and Chourey, 1999; Bihmidine et al., 2013). Based on the detection of early-endosperm E2F gene networks and their potential response to glucose-TOR pathway, we hypothesize that this pathway is involved in sensing the available post-phloem (in the vascular region of the pedicel and placento-chalazal region) photoassimilate levels and it either directly or indirectly signals the E2F gene networks to proliferate the developing coenocytic and cellular endosperm. In parallel, as endosperm cell types undergo differentiation during 3-4 DAP (during and after cellularization) (Leroux et al., 2014; Li et al., 2014), the formation of these cell types, including BETL, AL and ESR, likely impacts the rate of endosperm proliferation and the ultimate seed size. We detected signatures of a MYBR-regulated network likely driving BETL cell differentiation at 4 DAP. Because of its central role in transporting assimilates from the maternal tissue into the endosperm, proper BETL development is likely an important determinant of sink strength of the endosperm specifically and the seed in general (Ho, 1988; Olsen, 2004; Costa et al., 2012; Bihmidine et al., 2013; Haig, 2013). Our data suggests that a portion of the MYBR-regulated network shows altered gene expression in divergently selected lines of maize and may therefore be under artificial selection for seed size in cultivated maize. This provides a model in which the extent of BETL development is modulated in coordination with the overall rate of endosperm proliferation and its requirement for an optimum level of photoassimilates available in the early developing endosperm. Taken together, our identification of the gene networks associated with early endosperm proliferation and cell differentiation provide an entry point for analysis of underlying regulatory mechanisms that integrate physiological and developmental inputs such as sugar-sensing to control sink capacity and strength and ultimately determine seed size. This highlights the extent of the altered regulatory gene space necessary to modify seed size and points to specific gene networks as potential targets for its manipulation.

## Materials and Methods

### Plant materials

Maize (*Zea mays*) inbred line B73 plants were grown in a greenhouse with a cycle of 16 h of light at ∼30°C and 8 h of dark at ∼25°C at the University of Arizona during June-October 2012. Kernels at 0, 1, 2, 3, and 4 days after pollination (DAP) were harvested at the same time during the day (∼12 pm) from unfertilized (0 DAP) or self-pollinated (1-4 DAP) ears as previously described (Xin et al., 2013). Three biological replicates of each stage of endosperm development (0-4 DAP) and the embryo (4 DAP) were collected (18 samples total) for RNA profiling. The tissue sections for each biological replicate were pooled from at least eight individual kernels of a single ear (Supplemental Table 1). The *Nicotiana benthamiana* plants were grown in a growth chamber (Conviron) with a cycle of 16 h of light and 8 h of dark at 22°C. For mesophyll protoplast isolation, maize inbred line B73 plants were grown in the dark chamber ∼12 days at 25°C.

### Laser-capture microdissection, RNA isolation, cDNA amplification, RNA-Seq and data analysis

Laser-capture microdissection (LCM), RNA isolation and purification, cDNA amplification, and RNA-Seq were performed essentially as described previously (Zhan et al., 2015; Zhang et al., 2018). In RNA purification of the central cell and early endosperm stages (0-2 DAP), and the embryo (4 DAP), we repeatedly found indications of a carryover product(s) that caused an overestimation of RNA measurements using standard procedures. Instead, we used the height of the 18S-rRNA band from a Bioanalyzer 2100 (Agilent) analysis to measure RNA concentration. In all cases, ∼5 μg of amplified cDNA was obtained from ∼10 ng of captured RNA using the previously described amplification procedures (Zhan et al., 2015). The construction of the cDNA paired-end libraries and their quality checking were carried out using a TruSeq DNA Sample Preparation kit v2 (Illumina) with ∼1μg of amplified cDNA. Each bio-replicated set of cDNA samples was multiplexed and run on a single lane of an Illumina HiSeq 2500 system (Illumina) at the University of Arizona Genetics Core facility using the TruSeq SBS kit v3 (Illumina). Because of a failure of two samples within the first bio-replicate set, all samples for this bio-replicate were reprocessed from the library construction step through RNA-Seq, and the resulting total reads from both runs were combined for the subsequent analyses. Adapter sequences were trimmed from raw reads by the program Trimmomatic (Bolger et al., 2014) and sequence quality was checked with FastQC (Andrews, 2010).

Trimmed and high-quality reads were aligned to the maize reference genome (B73 RefGen_v3) using TopHat v2.0.9 (Trapnell et al., 2009). Intron length was set to 30 to 8000 nt with maximum number of mismatches per read set to 3 (-i 30 -I 8000 -N 3) (Zhan et al., 2015). Raw count table of exonic reads built by BEDTools v2.17.0 (Quinlan and Hall, 2010) with the exon coordinates (GTF, EnsemblPlants) were used in the software edgeR (Robinson et al., 2010), reported as counts per million mapped reads (CPM), and filtered with a lower limit of ≥0.5 in at least three of the 18 samples. The filtered raw count table was further normalized with the trimmed mean of M-values (TMM) method across the 18 samples, and transformed to CPM table using edgeR (Robinson et al., 2010). Annotation of the working gene set (WGS; 38,300 genes) and the high-confidence filtered gene set (FGS; 23,867 genes; focus of this study) were based on the v5b.60 gene models (B73 RefGen_v3, MaizeGDB).

### Identification of differentially expressed genes between endosperm and embryo

Two different comparison strategies with edgeR program (Robinson et al., 2010) were used to identify endosperm or embryo preferential genes in all FGS genes that were identified in our assay because the samples included five central cell/endosperm stages (collectively termed EN) and only one 4-DAP embryo (EMB) stage. For EN-pref genes determination, we first used a generalized linear model likelihood ratio test (3 bio-replicates, FC≥3, FDR ≤ 0.05) with two contrasts in comparison of group means (0-DAP + 1-DAP + 2-DAP + 3-DAP stage EN vs. 4 × EMB or 4-DAP EN vs. EMB) to identify EN upregulated genes, then applied the union to get the final gene list. For EMB-pref genes determination, we first used a generalized linear model likelihood ratio test (3 bio-replicates, FC ≤ 1/3, FDR ≤ 0.05) for a pairwise comparison of individual EN stage vs. EMB (0-DAP vs. EMB, 1-DAP vs. EMB, etc.) to identify EMB upregulated genes, then applied the intersection to get the final gene list.

### Identification of temporal co-expression modules using StepMiner

The StepMiner program (Sahoo et al., 2007) was used to identify genes with one-step-up, one-step-down, two-step-up-down, or two-step-down-up transition points from the expressed FGS gene set (23,867 genes) across central cell/endosperm time series using log_2_(CPM+1) values in all three bio replicates (*P* ≤ 0.05). The resulting 12,282 genes were further apportioned to the reported co-expression modules based on the stage of transition.

### *cis*-regulatory element analysis of temporal co-expression gene modules

MEME software (Bailey et al., 2006) was used to identify 10-12-bp-sequence motifs over-represented (top 20) within −1 kb to +0.5 kb (relative to the annotated transcription start site, TSS) of the genes in each StepMiner-identified co-expression module. Tomtom program (Gupta et al., 2007) was used to determine if any over-represented motifs were reported previously within the modified JASPAR CORE database (Mathelier et al., 2014) (q-value < 0.001). MRP-1-binding site (TATCTCTATCTC, bound by MRP-1) (Barrero et al., 2006) was manually added to the JASPAR CORE database. PatMatch (Yan et al., 2005) was used to detect occurrence of specific *cis*-motifs (Supplemental Table 10) upstream (−1 kb to +0.5 kb, relative to annotated TSS) of all genes in the genome. Individual *cis*-motif enrichment was determined for each temporal module using Fisher’s exact test (*P* < 0.001). Distribution of *cis*-motifs upstream (−1 kb to +0.5 kb, relative to annotated TSS) of all genes in the genome, or within one or more temporal module(s), was analyzed using a scaled density plot. The central position of the *cis*-motif locus was used to determine scaled densities.

### Gene functional enrichment analysis

Blast2GO (Conesa and Gotz, 2008) was used to perform Gene Ontology (GO) term enrichment analyses of the StepMiner-identified co-expression modules with a Fisher’s exact test (FDR < 0.05). GO annotations for maize genes were obtained from Gramene (gramene.org, Release 40). The enriched GO terms from each module were visualized using REVIGO (Supek et al., 2011).

### Phylogenetic analysis

Amino acid sequences were downloaded from EnsemblPlants (http://plants.ensembl.org). For genes with multiple isoforms, the longest one was used for the phylogenetic analyses. Sequences were aligned using Cluster Omega (EMBL-EBI; http://www.ebi.ac.uk), and a maximum likelihood phylogenetic tree was constructed using the Gamma model of rate heterogeneity and substitution matrix WAG in the RAxML (v. 7.7.1) web-server (Stamatakis et al., 2008). Bootstrap values were from 100 likelihood bootstrap replicates. Trees were visualized using FigTree (http://tree.bio.ed.ac.uk/software/figtree/).

### Yeast two-hybrid assays

The Matchmaker gold yeast two-hybrid (Y2H) system (Clontech) was used to identify protein-protein interactions among nine selected E2F/DP TFs. For each gene, the CDS was amplified with Phusion DNA polymerase (Thermo Fisher Scientific) from B73 endosperm cDNA using the primers described in Table S6 and cloned into the linearized vector pGADT7 (XmaI and BamHI) or pGBKT7 (EcoRI and BamHI) using In-Fusion HD cloning kit (Clontech). Y2H procedures were performed according to the Matchmaker gold yeast two-hybrid system user manual (Clontech). In brief, yeast *Saccharomyces cerevisiae* AH109 cells were co-transformed with the different bait (fused with a Gal4 DNA-binding domain, pGBKT7) and prey (fused with a Gal4-activation domain, pGADT7) combinations, and then grown on synthetic complete medium lacking Trp and Leu (double drop-out, DDO) and synthetic complete medium lacking Trp, Leu, His, and adenine (quadruple drop-out, QDO). Co-transformed colonies growing on QDO were considered as positive-interaction transformants.

### Dual luciferase reporter assays

Dual luciferase reporter (DLR) assay was used to detect binding and transactivation ability of E2Fs at the E2F-binding sites. The construct cloning and DLR assay of target-gene co-activation by two TFs were performed as previously described (Zhan et al., 2018) with minor modifications. In brief, for each gene, the CDS was amplified with Phusion DNA polymerase (Thermo Fisher Scientific) from B73 endosperm cDNA using the primers described in Table S7 and cloned into the linearized vector pBN-35Stev (XmaI and BamHI) using In-Fusion HD cloning kit (Clontech) to construct pBN-35Stev-TF. For each tested *cis*-motif, the DNA sequences of E2F-binding site concatemers with Gateway BP link were synthesized and cloned into the donor vector pDONR221 P3-P2 using Gateway BP reaction system (Table S7; Thermo Fisher Scientific). For multisite LR cloning, the entry clone pLAH-cismotif along with pLAH-Citrine, pLAH-VP64Ter, and the destination vector pLAH-LARm (Taylor-Teeples et al., 2015; Zhan et al., 2018) were used to generate the final expression construct pLAH-LARm-Citrine-VP64-cismotif. All clones were Sanger-sequenced to confirm the inserts. Leaves of *N. benthamiana* plants were co-infiltrated with a *A. tumefaciens* strain GV3101 (pMP90) harboring an expression construct (culture OD_600_ adjusted to 0.16), the p19 helper strain (culture OD_600_ adjusted to 0.1), and one or two pBN-35Stev-TF strain(s) (culture OD_600_ adjusted to 0.16). The absence of pBN-35Stev-TF strain in the co-infiltration was used as a control. Leaf tissue was collected, processed, and measured for firefly luciferase (LUC, as a reporter for transcriptional activation by the effector TF) vs the *Renilla* luciferase (REN, as a reporter for background signal correction)activities three days after infiltration. Five biological replicates were used for each set of assays. The pairwise comparisons of TF vs control or one TF vs two TFs were statistically tested using Student t-test (*P* < 0.05).

### Transient expression in protoplasts, RNA-Seq, and data analysis

For each gene, E2F CDS with nos terminator was amplified from pBN-d35Stev-E2F plasmid previously generated in DLR assay, cloned into EcoRV and BamHI linearized pCRII-TOPO-ubi vector (pCRII-TOPO vector containing ubiquitin promoter and 1st intron sequences) using In-Fusion cloning (Table S8; Clontech). For the following transient expression in protoplasts, highly pure and concentrated plasmids (>1 ug/ul) were isolated using ZymoPURE II plasmid midiprep kit (Zymo). The procedure of mesophyll protoplast isolation was modified from the published protocols (Yoo et al., 2007). The modifications included: The 2^nd^ leaves of 12 days old albino maize seedlings were used. Fifty leaves were used for protoplast collection. The middle part (∼7 cm) of leaves were chopped into 0.5 mm strips, and immediately transferred to 20 ml enzyme solution (0.6 M mannitol, 20 mM KCl, 20 mM MES, 1.5% Cellulase, 0.4% Macerozyme) in a Petri dish. The Petri dish was vacuumed for 30 min, and continued digestion for 4 h in the dark. Protoplasts were collected by passing through a 100 μm strainer (Falcon) with W5 solution (154 mM NaCl, 125 mM CaCl2, 5 mM KCl, and 2 mM MES, pH 5.7). The following steps of DNA-PEG-calcium transfection and protoplast harvest were exactly based on Yoo et. al (2007) protocol except 0.6 M mannitol used in WI and MMG solution, and 0.3 M mannitol in PEG solution. One or two plasmids with corresponding combinations were used for transfection (Dataset S13). After overnight (21 h) incubation of transfected protoplasts for transient expression in the dark, the protoplast samples were harvested for RNA-seq.

Total RNAs were isolated from protoplast samples using Direct-zol RNA kit (Zymo). The construction of the cDNA paired-end multiplexed sequencing libraries was carried out using the Illumina TruSeq Stranded mRNA Library Preparation Kit with poly(A) selection with 500 ng total RNA of each sample. The qualified and quantified libraries were pooled and sequenced on a single flow cell lane of the Illumina HiSeq 2500 instrument using the HiSeq 125 Cycle Paired-End Sequencing v4 protocol at the University of Utah High Throughput Genomics Core Facility.

RNA-seq data analysis procedures, including raw data preprocessing, quality-checking, mapping, read counting, were performed essentially as described previously (Zhan et al., 2018) with minor modification. In brief, trimmed reads were mapped to the maize reference genome (B73 RefGen_v3) using Tophat v2.1.1 (Trapnell et al., 2009). The raw count table was generated from mapped reads using featureCounts (Liao et al., 2014), and normalized with the TMM method across all samples and transformed to CPM table using edgeR (Robinson et al., 2010); the genes with raw counts per million > 1 in at least 3 of the 25 sequenced samples were used for above normalization. For differential expression analysis, DEGs were determined by pairwise comparisons (E2F(s) vs. control GFP) with a generalized linear model likelihood ratio test using edgeR (Robinson et al., 2010) under the cutoff criteria FDR < 0.05 with |log_2_FC| > log_2_(1.5). Collectively, the union set of E2F-upregulated DEGs further generated the E2F-up genes (Supplemental Figures 10A). Four bio-replicates were used for control, and three were used for all others.

### Electrophoretic mobility shift assays

Probe label and electrophoretic mobility shift assay (EMSA) were performed essentially as described previously (Zhan et al., 2018). The primers of constructions in E. coli expression were listed in Tables S9. All EMSA probes were labeled with Cy5 and were generated by annealing two Cy5-labeled oligonucleotides. The oligonucleotides used to generate the probes are listed in Tables S10. Unlabeled competitors were prepared by annealing oligonucleotides. The oligonucleotides used to generate the competitors are listed in Tables S10.

### Sugar-sensing, TOR-pathway-dependent gene analysis

Glc-TOR-pathway-dependent gene identification were extracted from the available data from Arabidopsis (Gene Expression Omnibus, GSE40245) (Xiong et al., 2013) using a filtering cutoff of *P*-value < 0.01 in both RMA and dChip analyses. Using EnsemblPlants, 3,374 Glc-TOR-up and 2,150 Glc-TOR-down maize homologs were identified in our FGS dataset as outlined in Supplemental Figure 11B. A list of 1,021 candidate KIN10 target genes based on transient KIN10 expression in Arabidopsis protoplasts (Baena-Gonzalez et al., 2007) was used to obtain the maize homologs using EnsemblPlants. A total of 601 KIN10-up and 734 KIN10-down maize homologs were identified in our FGS dataset (Supplemental Figure 11E). Associations of Glc-TOR- and KIN10-regulated genes with the StepMiner-identified co-expression modules in maize endosperm were determined using Fisher’s exact test (*P* < 0.001).

### Gene annotation and analysis of functional categories

For all FGS genes, the corresponding Zm-B73-REFERENCE-GRAMENE-4.0 Gene ID (Gene ID_v4) and Zm-B73-REFERENCE-NAM-5.0 Gene ID (Gene ID_v5) were converted in MaizeGDB(Lawrence et al., 2004citations). Gene descriptions in maize were obtained from EnsemblPlants. Locus names were downloaded from MaizeGDB (Lawrence et al., 2004citations)). Locus names mentioned here were manually modified based on the references, including those for cyclins (Hu et al., 2010) and trehalose-6-phosphate synthases and phosphatases (Henry et al., 2014). Annotation of TF family members were based on data from Plant TF database v3.0 (PlantTFDB 3.0) (Jin et al., 2014) and GrassTFDB in GRASSIUS (Yilmaz et al., 2009). Purα genes GRMZM2G049429 and GRMZM2G069208 were manually added to the Orphans family of TFs based on previously published data (Tremousaygue et al., 1999; Tremousaygue et al., 2003; Johnson et al., 2013). Analysis of the gene functional categories was based on the MapMan annotation data (Thimm et al., 2004) and a previous report (Xiong et al., 2013). *P*-values of functional enrichments were calculated using Fisher’s exact test.

### Statistical analysis

All plots were constructed in R with data presented as mean of the three bio-replicates unless otherwise noted. Spearman correlation coefficient (SCC) was used to determine the extent of reproducibility of mRNA populations between the three bio-replicates of each stage. SCC was calculated from log2-transformed CPM (i.e., log_2_(CPM+1)) using the cor.test function. The relationship among the mRNA populations of the 18 samples was plotted with a multidimensional scaling (MDS) based on Euclidean distances of normalized expression level (log_2_(CPM+1)) using edgeR (Robinson et al., 2010). All determinations of under- or over-representation of genes in a given category were based on Fisher’s exact test. Wilcoxon rank-sum test was used to determine the differential distributions of the large- vs. small-sized-kernel gene expression in temporal modules or subsets thereof using normalized RNA reads (in FPKM) for endosperm and embryo RNA profiles obtained from two divergently selected maize populations of large (KLS30) and small (KSS30) kernels (Sekhon et al., 2014).

## Acknowledgments

We thank Chuang Ma and Ruolin Yang for advice on the use of bioinformatics tools, Haijiao Wang for help with tissue collection and fixation, Alan Lloyd for making EMSA constructs and doing EMSA experiments, Samuel Hazen for providing the pLAH-VP64Ter and pLAH-LARm vectors, Guang Yao for providing the pDONR221-P1-P4 and pDONR221-P3-P2 vectors, Wang Tian and Donna D. Zhang for providing instrument supports of the GloMax 20/20 Luminometer, and Mark Beilstein for advice on the phylogenetic analysis. We also thank Karen Schumaker for critical reading of the manuscript. This work was supported by the National Science Foundation grants IOS-0923880 and IOS-1444568 (to J.M.D., G.N.D. and R.Y.).

## Author Contributions

S.Z., J.M.D., G.N.D., and R.Y. designed the experiments; S.Z., C.-H.R., and G.L. performed the experiments; S.Z., D.R., X.W. and R.Y. performed data analysis; and S.Z. and R.Y. wrote the manuscript.

## Data availability

The RNA-sequencing data have been deposited in the Gene Expression Omnibus (GEO) under accession GSE69784.

## Disclosure Declaration

The authors declare no conflict of interest.

## Notes

### Competing Interest Statement

The authors have declared no competing interest.

