## Supplementary figures and images for "Gene networks associated with early endosperm proliferation and basal endosperm layer differentiation in maize"

### Supplemental Figure 1

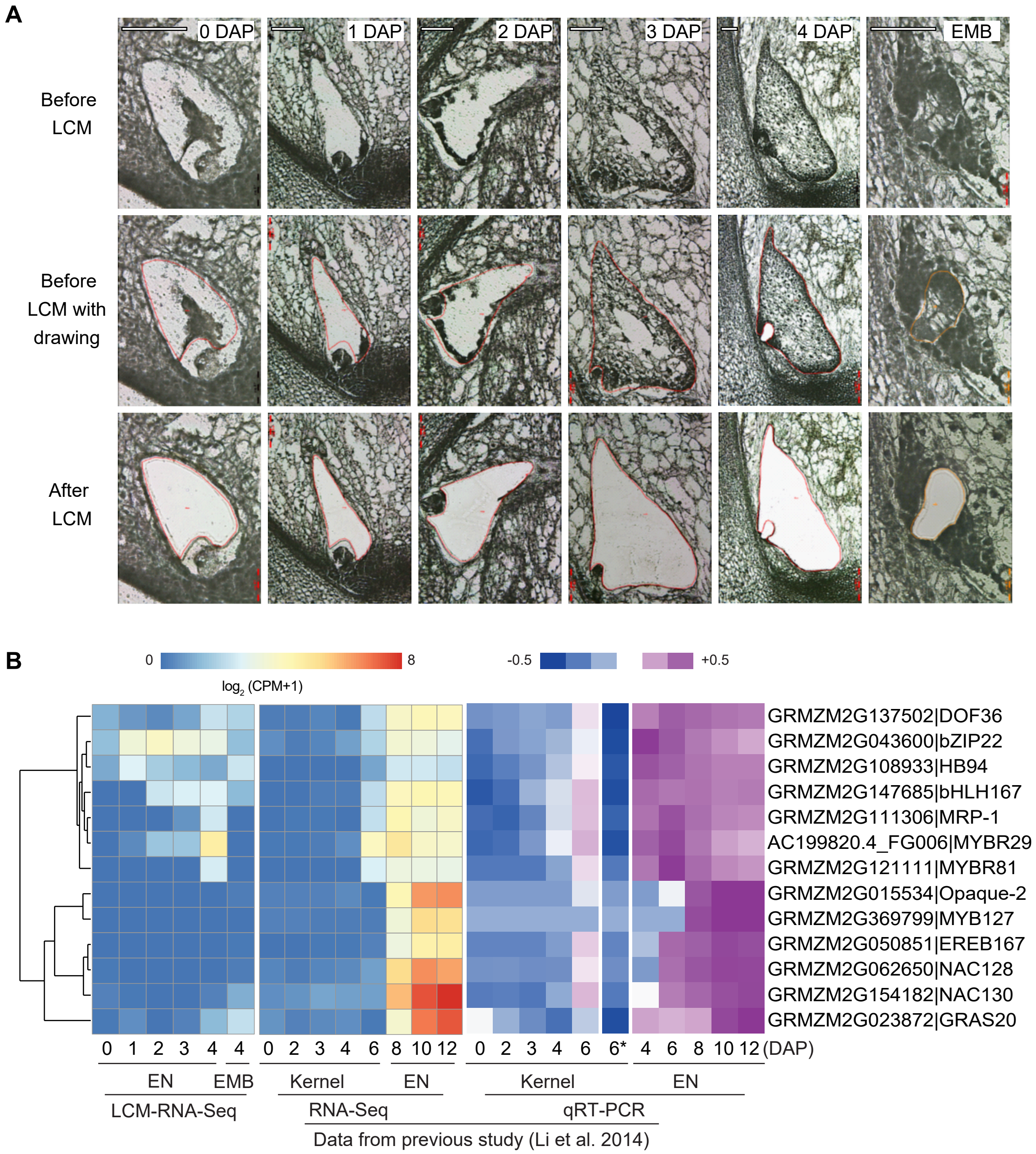


**Supplemental Figure 1**

### Supplemental Figure 2

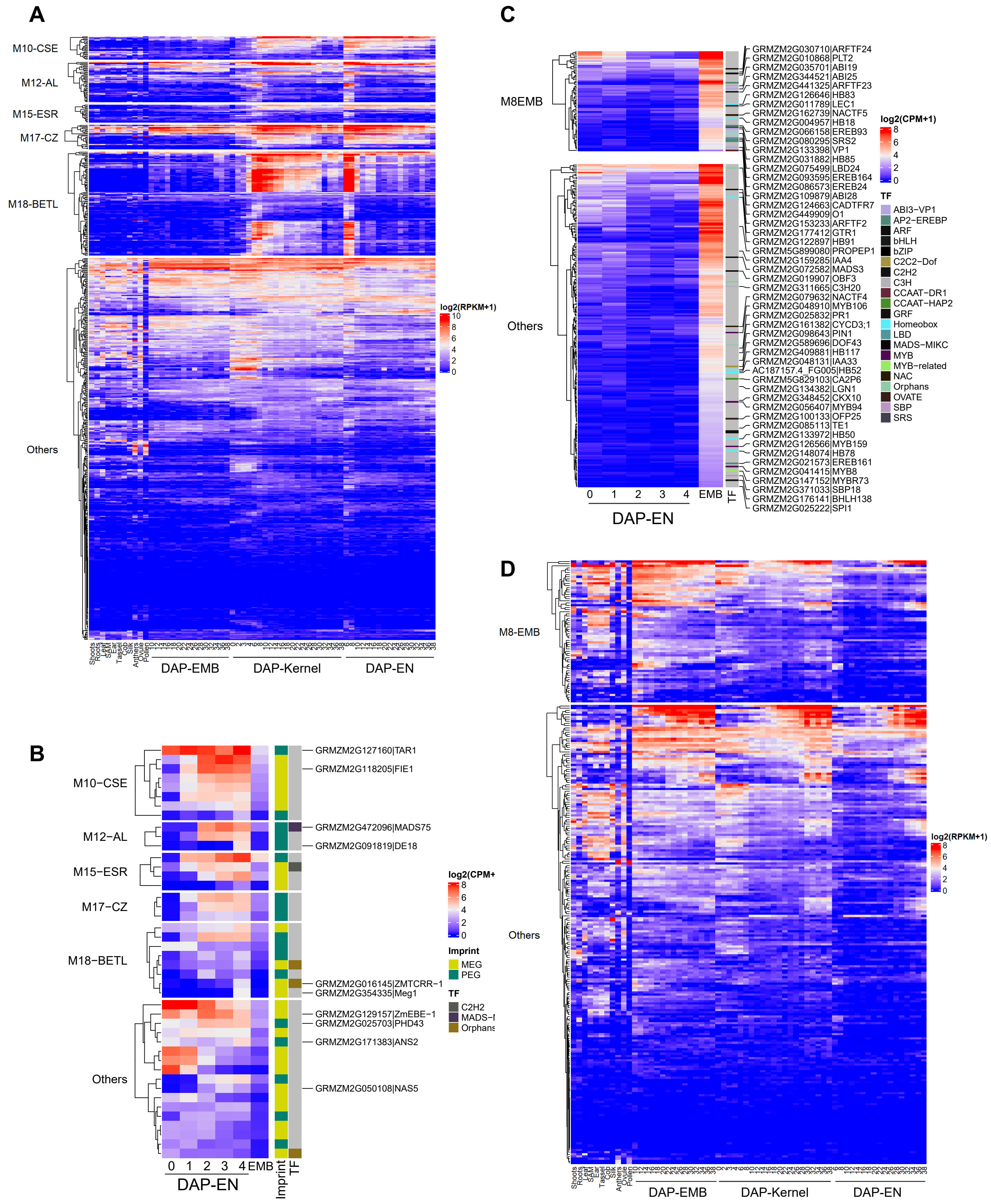


**Supplemental Figure S2**

### Supplemental Figure 3

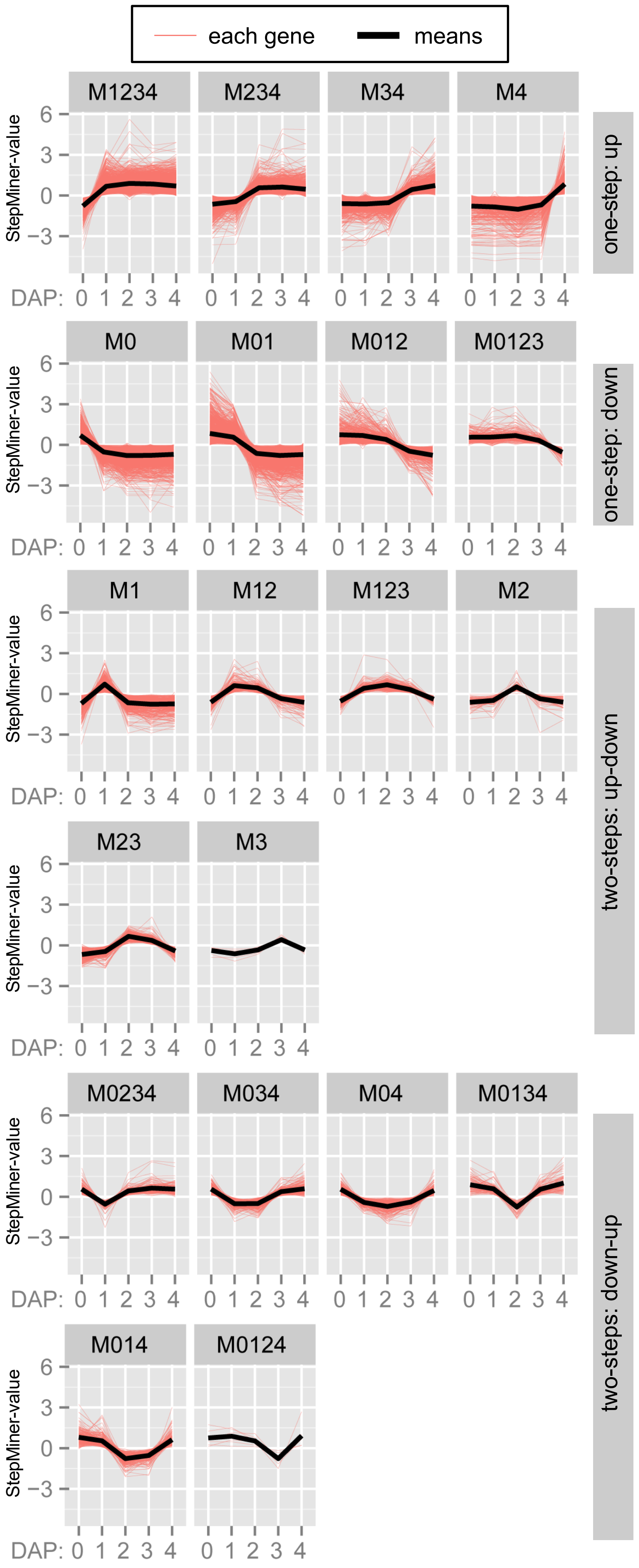


**Supplemental Figure 3**

### Supplemental Figure 4

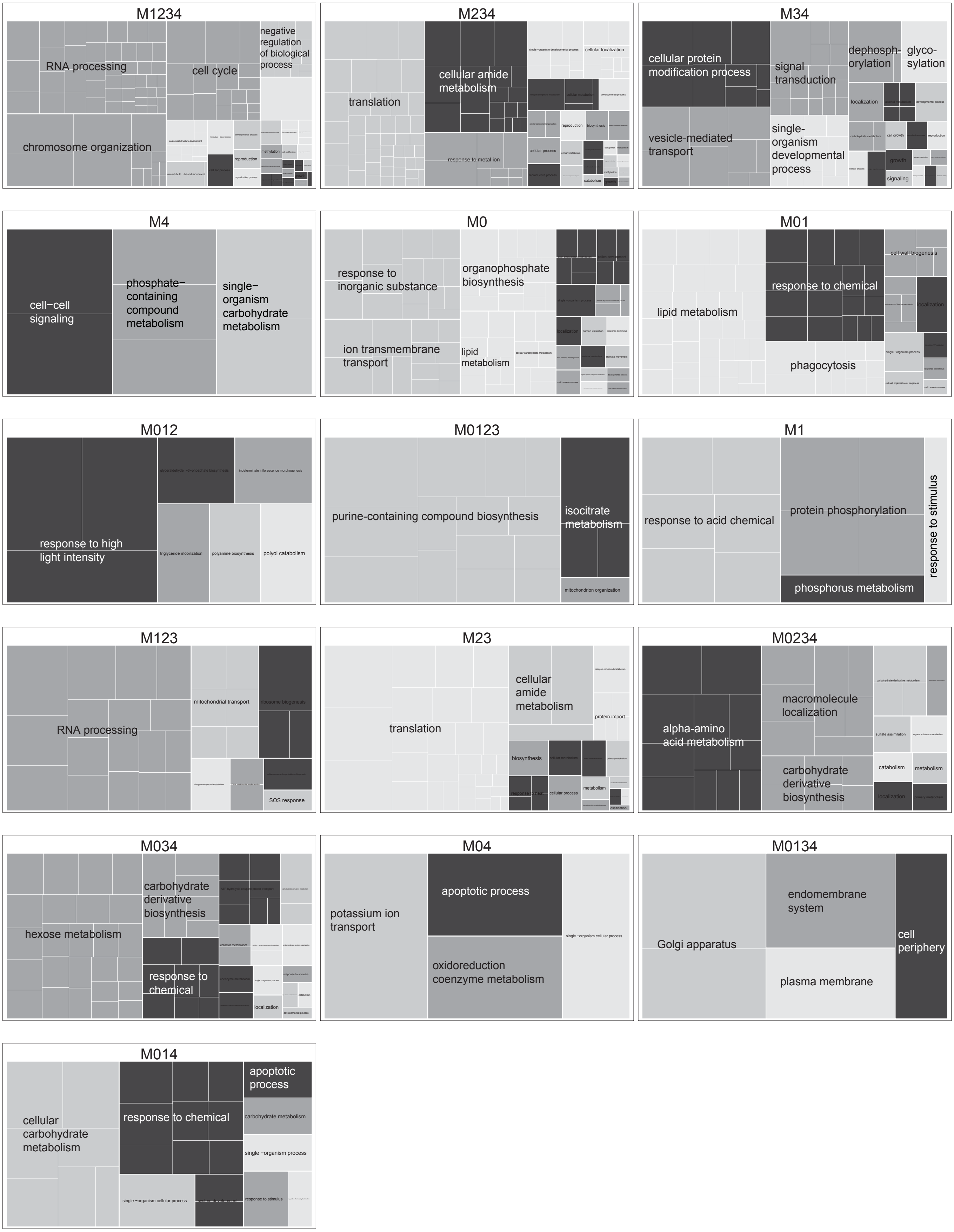


**Supplementary Figure 4**

### Supplemental Figure 5

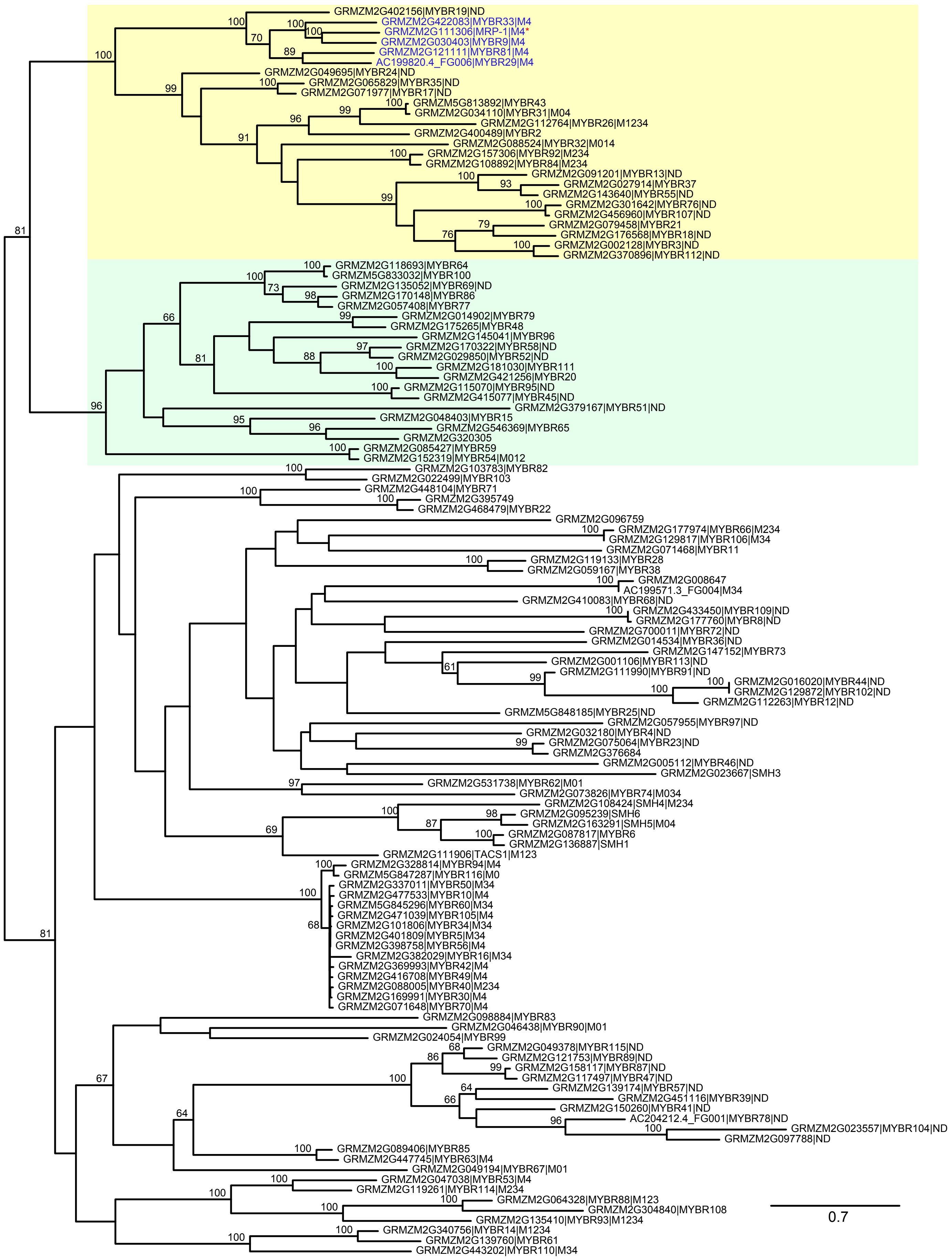

**Supplemental Figure 5**

### Supplemental Figure 6

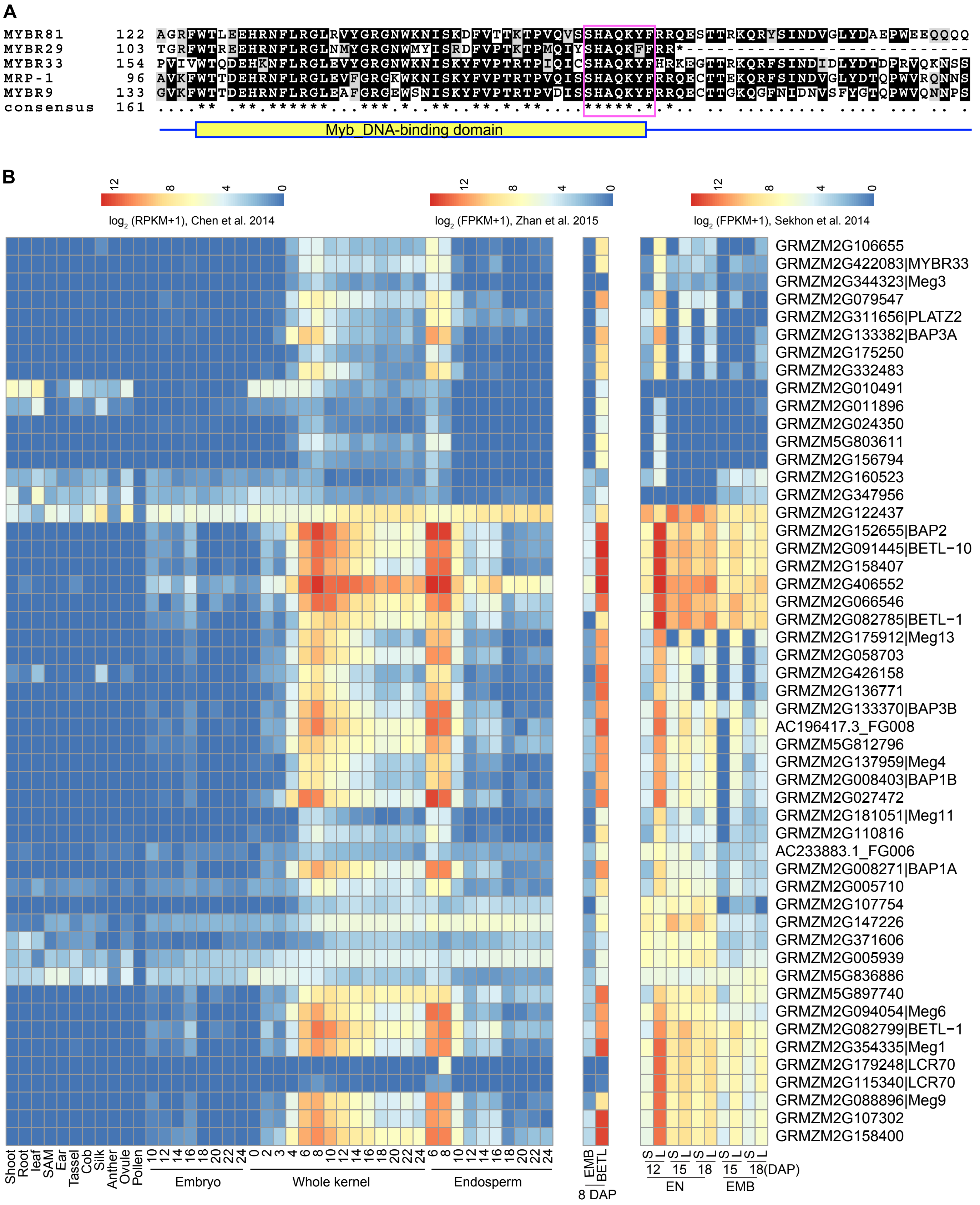


**Supplemental Figure 6**

### Supplemental Figure 7

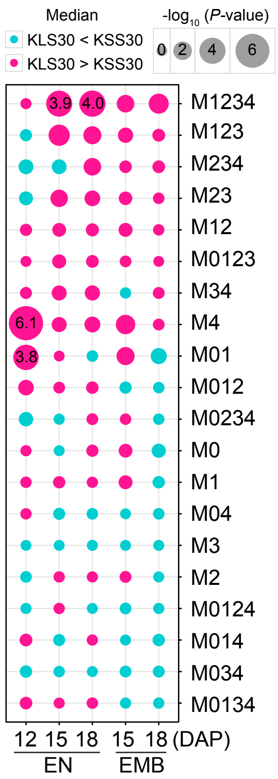


**Supplemental Figure 7**

### Supplemental Figure 8

**A**

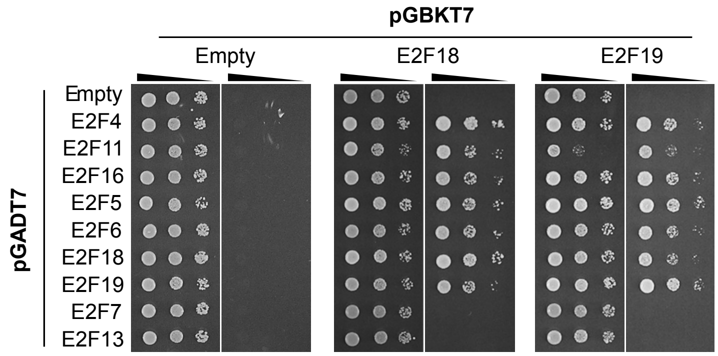


**B**

| | BD | E2F5 | E2F6 | E2F18 | E2F19 | E2F7 |
|---|---|---|---|---|---|---|
| AD | - | - | - | - | - | - |
| E2F4 | - | + | + | + | + | - |
| E2F11 | - | + | + | + | + | - |
| E2F16 | - | + | + | + | + | - |
| E2F5 | - | + | + | + | + | - |
| E2F6 | - | + | + | + | + | - |
| E2F18 | - | + | + | + | + | - |
| E2F19 | - | + | + | + | + | - |
| E2F7 | - | - | - | - | - | - |
| E2F13 | - | - | - | - | - | - |

**C**

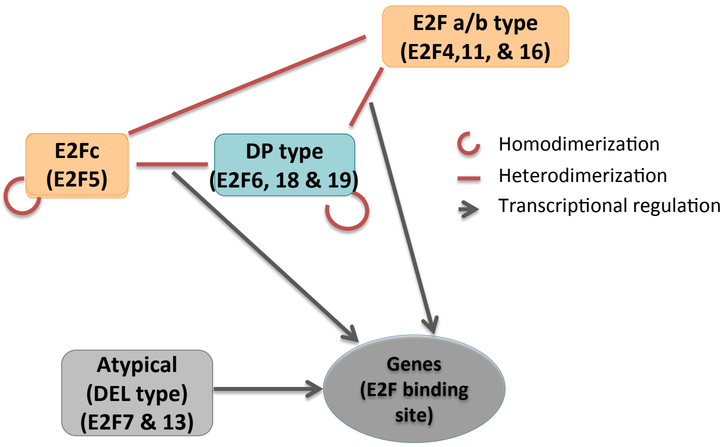


**Supplemental Figure 8**

### Supplemental Figure 11

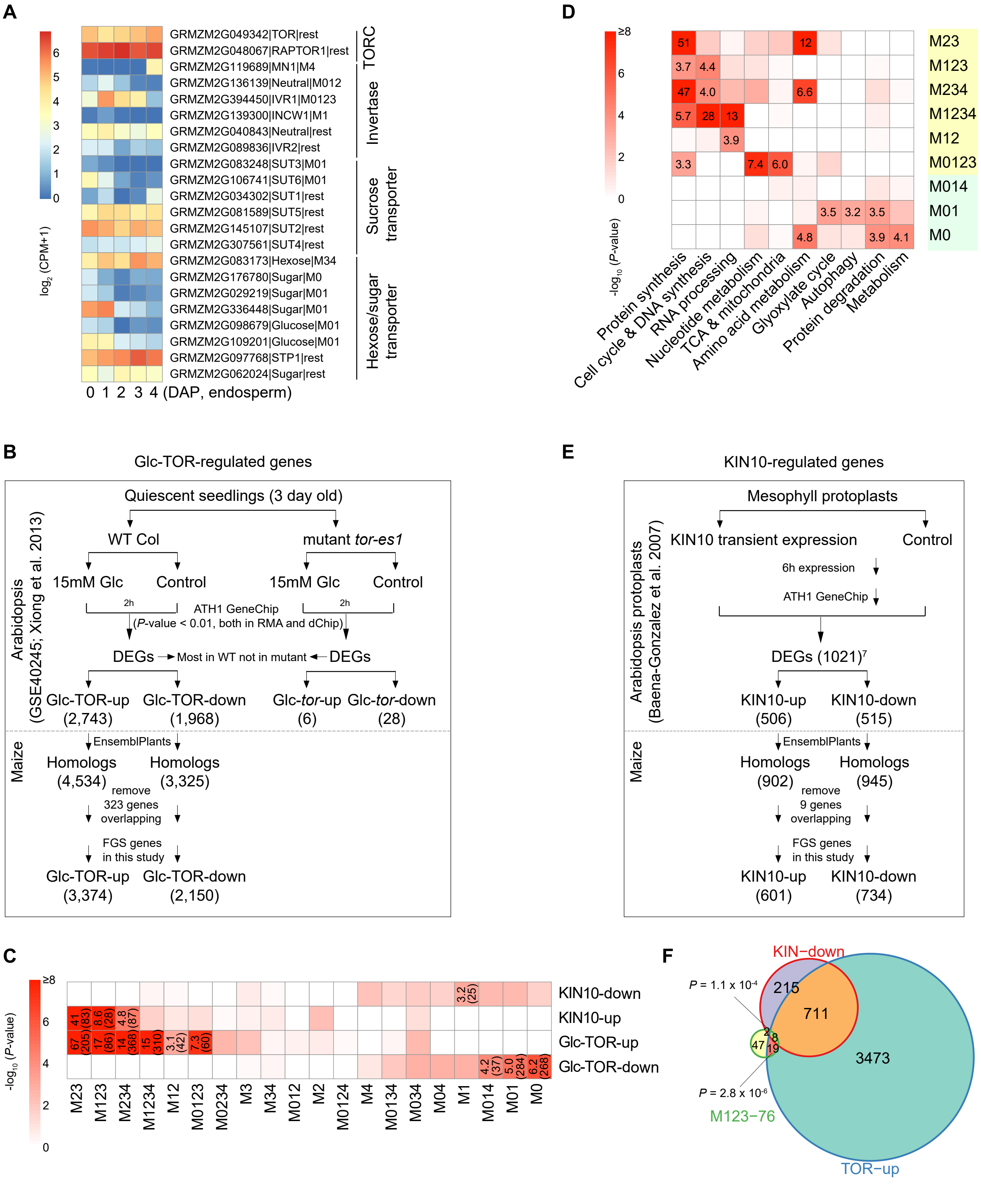


**Supplemental Figure 11**
