## Supplemental Figure 9 for "Gene networks associated with early endosperm proliferation and basal endosperm layer differentiation in maize"

**A**

| Submotif | Sequences | M1234 | | M123 | |
|---|---|---|---|---|---|
| | | Gene # | % Gene | Gene # | % Gene |
| a | TCTCCCGC | 160 | 46.9 | 38 | 50.0 |
| b | TTTCCCGC | 76 | 22.3 | 12 | 15.8 |
| c | TTTGGCGC | 70 | 20.5 | 14 | 18.4 |
| d | TTTGCCGC | 60 | 17.6 | 16 | 21.1 |
| e | TTTCGCGC | 37 | 10.9 | 2 | 2.6 |

**B**

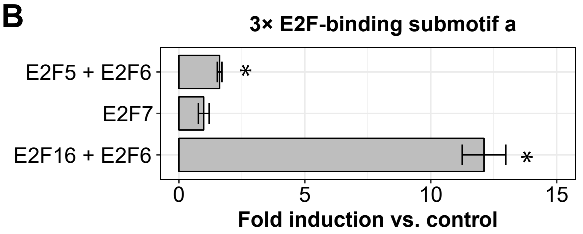


**C**

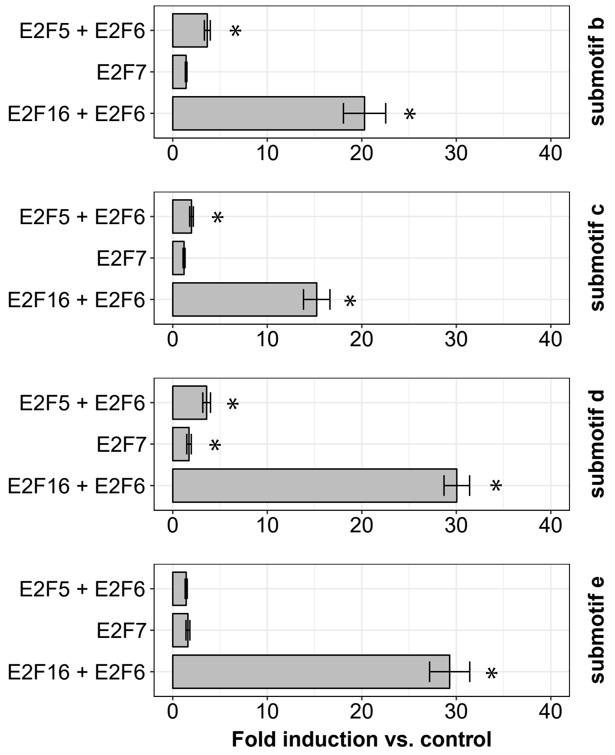


**D**

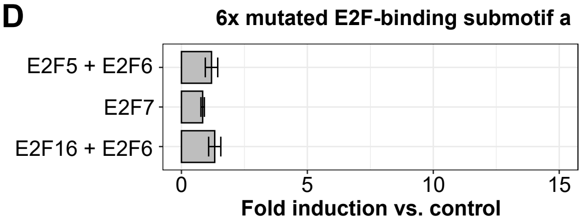


**E**

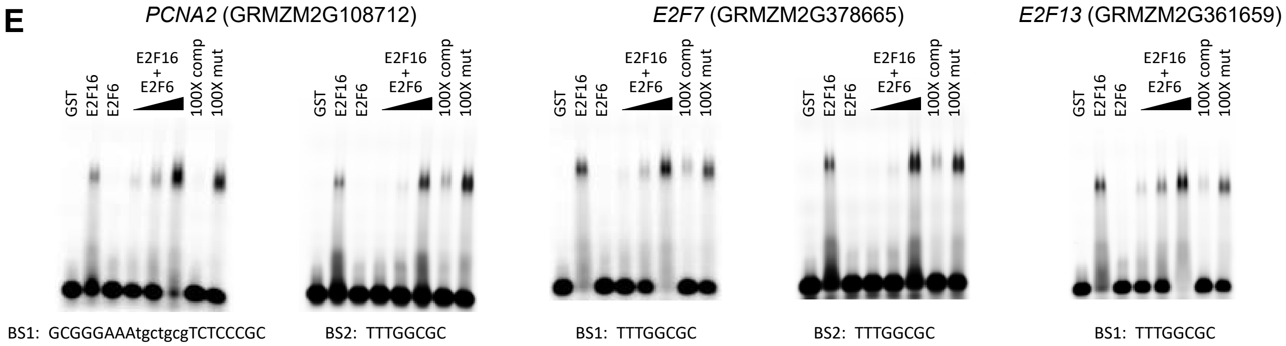


**Supplemental Figure 9**
