## Supplemental Figure 10 for "Gene networks associated with early endosperm proliferation and basal endosperm layer differentiation in maize"

**A**

| Pairwise comparison | DEG # Up | DEG # Down | Name of combined DEG |
|---|---|---|---|
| E2F5 vs GFP | 16 | 12 | E2F5 DEGs |
| E2F5 & E2F6 vs GFP | 135 | 18 | |
| E2F6 vs GFP | 2 | 4 | |
| E2F7 vs GFP | 1316 | 720 | E2F7 DEGs |
| E2F16 vs GFP | 3444 | 2494 | E2F16 DEGs |
| E2F16 & E2F6 vs GFP | 4417 | 2485 | |
| Union | 6163 | - | E2F-up genes |

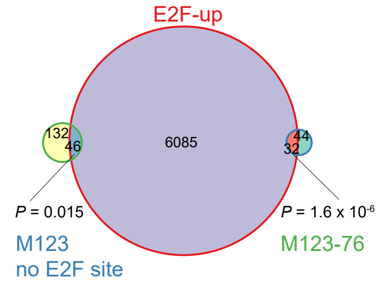


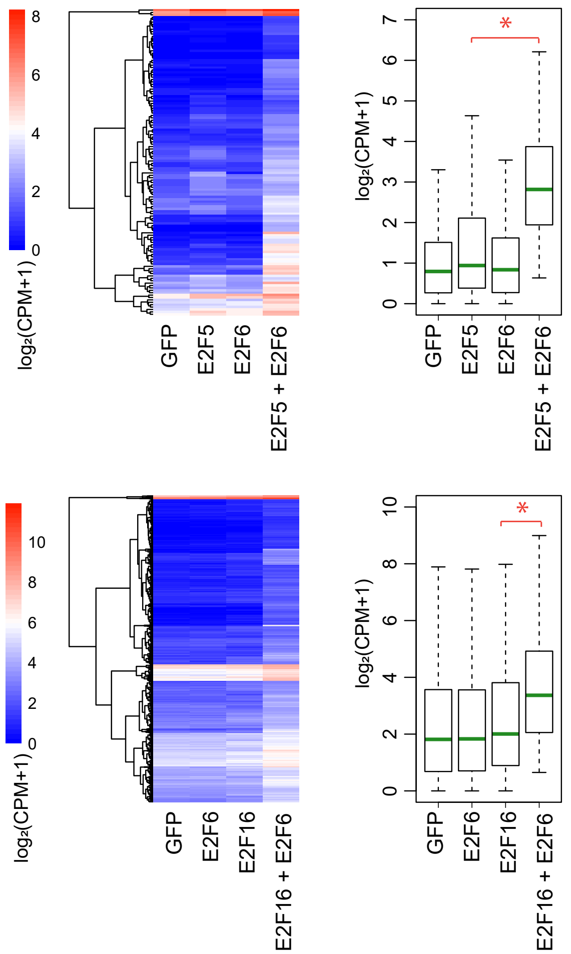


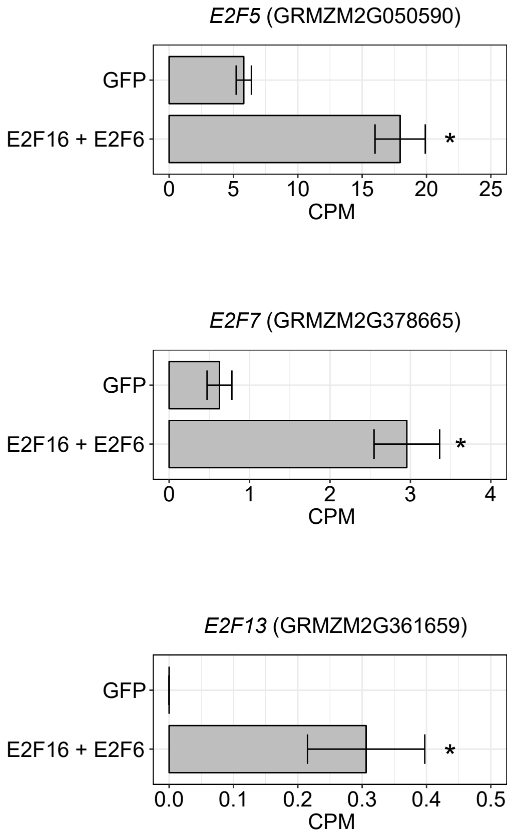


**Supplemental Figure 10**
