## Supplemental Table 1 for "Gene networks associated with early endosperm proliferation and basal endosperm layer differentiation in maize"

**Table S1** LCM sampling and RNA yields.

| Samples^a^ | # kernels | Captured area (mm^2^)^b^ | RNA yield (ng) |
| --- | --- | --- | --- |
| Endosperm, 0 DAP, rep-1 (0 DAP-1) | 87 | 3.47 | 37.2 |
| Endosperm, 0 DAP, rep-2 (0 DAP-2) | 84 | 3.72 | 49.5 |
| Endosperm, 0 DAP, rep-3 (0 DAP-3) | 95 | 4.47 | 54.6 |
| Endosperm, 1 DAP, rep-1 (1 DAP-1) | 51 | 2.96 | 32.3 |
| Endosperm, 1 DAP, rep-2 (1 DAP-2) | 45 | 2.80 | 40.0 |
| Endosperm, 1 DAP, rep-3 (1 DAP-3) | 47 | 3.04 | 42.3 |
| Endosperm, 2 DAP, rep-1 (2 DAP-1) | 16 | 3.44 | 66.8 |
| Endosperm, 2 DAP, rep-2 (2 DAP-2) | 18 | 5.24 | 102.8 |
| Endosperm, 2 DAP, rep-3 (2 DAP-3) | 19 | 4.11 | 121.5 |
| Endosperm, 3 DAP, rep-1 (3 DAP-1) | 17 | 35.01 | 477.7 |
| Endosperm, 3 DAP, rep-2 (3 DAP-2) | 9 | 8.67 | 267.2 |
| Endosperm, 3 DAP, rep-3 (3 DAP-3) | 11 | 10.71 | 169.1 |
| Endosperm, 4 DAP, rep-1 (4 DAP-1) | 17 | 100.00 | 2946.0 |
| Endosperm, 4 DAP, rep-2 (4 DAP-2) | 8 | 37.78 | 1275.2 |
| Endosperm, 4 DAP, rep-3 (4 DAP-3) | 8 | 29.89 | 1133.1 |
| Embryo, 4 DAP, rep-1 (EMB-1) | 17 | 0.36 | 50.3 |
| Embryo, 4 DAP, rep-2 (EMB-2) | 19 | 0.24 | 31.7 |
| Embryo, 4 DAP, rep-3 (EMB-3) | 15 | 0.36 | 29.9 |

^a^A developmental series of the endosperm at 0, 1, 2, 3, and 4 DAP, and the whole embryo at 4 DAP with three bio-replicates (rep) in maize inbred B73. Each bio-replicate was obtained from an individual ear. DAP: Day after pollination. ^b^Section thickness = 10 µm. When slides were prepared, some sections from a kernel might be lost as the ribbons were broken. Therefore, not all structures in some sections (from a single kernel) might be fully captured if they were out-of-frame or not clearly discernible.
