## Supplemental Table 2 for "Gene networks associated with early endosperm proliferation and basal endosperm layer differentiation in maize"

**Supplemental Table 2** Reads and mapping statistics of LCM-RNA-Seq data.

| Samples | Total # reads | All mapped read | | Mapped to exons | |
| --- | --- | --- | --- | --- | --- |
|  |  | # | % | # | % (of mapped) |
| 0 DAP-1 | 91,274,932 | 70,801,895 | 77.6 | 19,395,626 | 27.4 |
| 0 DAP-2 | 77,470,152 | 60,517,493 | 78.1 | 20,655,583 | 34.1 |
| 0 DAP-3 | 52,156,636 | 42,884,009 | 82.2 | 10,684,194 | 24.9 |
| 1 DAP-1 | 14,052,010 | 9,176,962 | 65.3 | 2,096,112 | 22.8 |
| 1 DAP-2 | 66,364,646 | 52,286,590 | 78.8 | 16,613,378 | 31.8 |
| 1 DAP-3 | 51,585,100 | 41,646,677 | 80.7 | 13,181,322 | 31.7 |
| 2 DAP-1 | 97,050,066 | 72,736,995 | 74.9 | 29,964,707 | 41.2 |
| 2 DAP-2 | 44,582,200 | 37,385,515 | 83.9 | 19,081,924 | 51.0 |
| 2 DAP-3 | 61,258,024 | 51,079,405 | 83.4 | 24,851,116 | 48.7 |
| 3 DAP-1 | 56,395,412 | 34,605,131 | 61.4 | 16,062,501 | 46.4 |
| 3 DAP-2 | 24,008,794 | 19,884,689 | 82.8 | 10,180,970 | 51.2 |
| 3 DAP-3 | 55,303,094 | 45,722,431 | 82.7 | 23,363,196 | 51.1 |
| 4 DAP-1 | 85,818,772 | 58,053,484 | 67.6 | 24,947,377 | 43.0 |
| 4 DAP-2 | 43,262,606 | 35,233,226 | 81.4 | 16,442,357 | 46.7 |
| 4 DAP-3 | 49,134,140 | 40,614,926 | 82.7 | 18,209,578 | 44.8 |
| EMB-1 | 16,847,222 | 10,361,188 | 61.5 | 2,497,803 | 24.1 |
| EMB-2 | 65,766,532 | 50,907,306 | 77.4 | 19,087,621 | 37.5 |
| EMB-3 | 43,186,350 | 35,040,288 | 81.1 | 12,087,566 | 34.5 |
| Total | 995,516,688 | 768,938,210 | 77.2 | 299,402,931 | 38.9 |
