## Supplemental Table 3 for "Gene networks associated with early endosperm proliferation and basal endosperm layer differentiation in maize"

**Table S3** Spearman correlation coefficient analysis of the replicates of the LCM-RNA-Seq libraries.

| Stages | rep-1 vs. rep-2 | rep-1 vs. rep-3 | rep-2 vs. rep-3 |
| --- | --- | --- | --- |
| 0 DAP | 0.94 | 0.91 | 0.95 |
| 1 DAP | 0.91 | 0.90 | 0.96 |
| 2 DAP | 0.95 | 0.95 | 0.96 |
| 3 DAP | 0.94 | 0.94 | 0.97 |
| 4 DAP | 0.95 | 0.95 | 0.98 |
| EMB | 0.89 | 0.90 | 0.95 |
