## Supplemental Table 4 for "Gene networks associated with early endosperm proliferation and basal endosperm layer differentiation in maize"

**Table S4** Strategies of preferentially expressed gene identification in endosperm and embryo.

| **Contrast** | **FC^a^** | **Gene #** | **Union/Intersection^b^** | | |
| --- | --- | --- | --- | --- | --- |
|  |  |  | **Gene #** | **TF #** | **Term** |
| 0-DAP + 1-DAP + 2-DAP + 3-DAP stage EN vs. 4 × EMB | ≥ 3 | 319 | 588 | 47 | EN-pref |
| 4-DAP EN vs. EMB | ≥ 3 | 350 |  |  |  |
| 0-DAP EN vs. EMB | ≤ 1/3 | 1180 | 275 | 46 | EMB-pref |
| 1-DAP EN vs. EMB | ≤ 1/3 | 718 |  |  |  |
| 2-DAP EN vs. EMB | ≤ 1/3 | 807 |  |  |  |
| 3-DAP EN vs. EMB | ≤ 1/3 | 759 |  |  |  |
| 4-DAP EN vs. EMB | ≤ 1/3 | 730 |  |  |  |

^a^Using edgeR with GLM likelihood ratio tests, 3 bio-replicates, FDR ≤ 0.05 with corresponding cutoff on Fold Change (FC). ^b^EN-pref gene set was determined by union, and EMB-pref gene set was determined by intersection.
