## Supplemental Table 5 for "Gene networks associated with early endosperm proliferation and basal endosperm layer differentiation in maize"

**Table S5** *cis*-motifs used for PatMatch analysis of the temporal modules.

| ***cis*-motif** | **Alternative name** | **PatMatch analysis** | | **Reported TF regulators** | | **Reported genes possessing *cis*-motif** | | **Refs** |
| --- | --- | --- | --- | --- | --- | --- | --- | --- |
|  |  | **Motif sequence^a^** | **Enriched modules^b^** | **Associated TFs** | **Species** | **Characteristics (e.g.)** | **List (e.g.)^c^** |  |
| *telo* box | Short interstitial telomere motif | AAACCCTA^1^ | M1234, M234, M12, M123, M23 | Purα | Arabidopsis | Components of the translational machinery | *EF1α, RPGs, snRNP, OsRPGs, OssnRNP* | 1-4 |
| E2F-binding sites | E2F element | TTTSSCGC{Naouar, 2009 #111}/ | M1234, M123 | E2F | Arabidopsis, maize, rice, tobacco | Expressed during G1 and S phases | *PCNA, RNR, MSH, CYCA, CYCB1;2, ORC1, OsPCNA, NtPCNA, ZmRBR3* | 7-13 |
|  |  | TCTCCCGC^11^ |  |  |  |  |  |  |
| MRP-1-binding sites | GATA-rich sequence | TAGATAGATAGA/ | M4 | MRP-1 (maize) | maize | Expressed in the basal endosperm transfer layer | *Meg1, BAP, BETL (*all maize*)* | 19-21 |
|  |  | TAGATATAGATA/ |  |  |  |  |  |  |
|  |  | TAGATAGAGATA/ |  |  |  |  |  |  |
|  |  | TAGATAGATATA^21^ |  |  |  |  |  |  |

^a^H = A, C, or T; R = A or G; Y = C or T; W = A or T. ^b^*P*-values were calculated by Fisher’s exact test. *P* < 0.001. ^c^RPGs: Ribosomal protein genes. Genes from rice, tobacco, and maize are prefixed Os, Nt, and Zm, respectively. Genes without any note are from Arabidopsis.
