## Supplemental Information for "Gene networks associated with early endosperm proliferation and basal endosperm layer differentiation in maize"

**TITTLE:**

NOTES:

^1^ Current address: KWS Group, KWS Gateway Research Center, 1005 N Warson Rd, Saint Louis, MO 63132, USA.

^2^ Current address: Department of Plant Genetics and Breeding, China Agricultural University, Beijing 100193 China

**SUPPLEMENTAL INFORMATION**

**SUPPLEMENTAL FIGURE LEGENEDS**

**Supplemental Figure 1.** Laser-capture microdissection of early endosperm and embryo, and validation of the captured RNAs. (A) A representative sample of paraffin sections of kernels at 0, 1, 2, 3 and 4 DAP before, during, and after laser-capture microdissection (LCM). The area selected for LCM is outlined in red (endosperm) or orange (embryo). Bars, 100 µm. (B) Analysis of mRNA levels of 13 transcription-factor (TF) genes with data from this study (LCM-RNA-Seq time series) and a previous study (RNA-Seq and qRT-PCR datasets) (Li et al., 2014) using a hierarchically clustered heat map. From the latter study, RNA-Seq data were re-analyzed and transformed to CPM by edgeR (Robinson et al., 2010) while normalized Ct values of qRT-PCR assays were used directly here. 6* denotes 6-DAP kernels with the endosperm removed (Li et al., 2014).

**Supplemental Figure 2.**  Identification of endosperm- vs. embryo-preferentially expressed genes in our LCM-RNA-Seq study. (A) Relative expression levels of EN-pref gene sets visualized using a hierarchically clustered heat map based on normalized RNA reads in RPKM from the maize B73 expression atlas (Chen et al., 2014) with staged organs and tissues including shoot, root, leaf 5, shoot apical meristem (SAM 1), ear 1, tassel 4, pre-emergence cob 1, silk, anther, ovule, pollen, whole kernels, endosperm, and embryos of different time-series stages (in DAP). Genes were initially clustered based on previously described gene co-expression modules obtained from 8-DAP kernels (Zhan et al., 2015). (B) Relative expression levels of the imprinted genes in EN-pref gene sets visualized using a hierarchically clustered heat map based on normalized RNA reads in CPM in our LCM transcriptome data. The genes were initially clustered as described for (A). The right bar annotations of heat map include the imprinting patterns and TF family information. The genes with publicly given names are reported on the right. Maternally expressed genes (MEGs) and paternally expressed genes (PEGs) are color coded. Select genes are noted. All associated TF gene families are identified by color-coding. (C) Relative expression levels of EMB-pref gene sets visualized using a hierarchically clustered heat map based on normalized RNA reads from our LCM-RNA-Seq data and grouped as in (A). All associated TF gene families are identified by color-coding. The genes with publicly given names are reported on the right. (D) Relative expression levels of EMB-pref gene sets visualized using a hierarchically clustered heat map based on normalized RNA reads in RPKM from the expression atlas of maize B73 (Chen et al., 2014). The staged organs and tissues are as in (A).

**Supplemental Figure 3.**  Additional analyses of temporal co-expression modules. Temporal mRNA accumulation profiles of the 20 StepMiner-identified co-expression modules. Profiles of individual genes are shown as red lines, and average profiles of all genes in each module are shown as black lines.

**Supplemental Figure 4.**  Enrichment of GO terms for temporal co-expression modules analyzed using Blast2go (FDR < 0.05) and visualized using ReviGO treemap with FDR values. All supporting details including GO terms and FDR values are indicated in Dataset S16.

**Supplemental Figure 5.**  Phylogenetic tree of MYBR TF family in maize using predicted amino acid sequences. All 25 proteins in the phylogenetic clade highlighted in yellow and 13 of 20 proteins in the clade highlighted in green have a highly conserved SHAQK(Y/F)F amino acid domain. The proteins containing the SHAQK(Y/F)F domain in the M4 module are indicated in blue font. Where applicable, the temporal module expression pattern associated with each maize gene is indicated as an extension of the gene name. ND denotes not detected based on our expression datasets. The maximum likelihood tree was constructed using the program RAxML (Stamatakis et al., 2008). Values above nodes are from 100 bootstrap replicates; only values >60% are reported. Bar, 0.7 amino acid substitutions per site.

**Supplemental Figure 6.**  Candidate MYBR TF proteins associated with the M4-47 temporal module subset and global analysis of their mRNA levels using the available expression data. (A) Alignment of M4-47 candidate MYBR proteins over the Myb DNA-binding domain region showing a highly conserved SHAQK(Y/F)F amino acid sequence signature. Prediction of the Myb DNA-binding domain was based on the available data in PlantTFDB 3.0 (Jin et al., 2014). (B) Relative levels of M4-47 gene mRNAs visualized using a heat map based on normalized RNA reads for select tissues/organs from three published RNA-Seq datasets including an expression atlas of maize inbred B73 with shoot, root, leaf 5, shoot apical meristem (SAM 1), ear 1, tassel 4, pre-emergence cob 1, silk, anther, ovule, pollen, whole kernels, endosperm, and embryos of different developmental stages (in DAP) (Chen et al., 2014) (RPKM, left), a BETL and embryo (EMB) LCM profile of 8-DAP kernel compartments (Zhan et al., 2015) (FPKM, middle), and a profile of endosperm and embryo RNAs obtained from two divergently selected maize populations of large (KLS30) and small (KSS30) sized kernels (Sekhon et al., 2014) (FPKM, right). The expression of most M4-47 genes was extremely low in embryos compared to endosperm except in the latter dataset. Based on these profiles, we conclude that the differences in mRNA levels obtained from 15- and 18-DAP embryos of large vs. small seeds (Figure 3D) are likely due to artifacts associated with mRNA preparations (*i.e.,* contamination of embryos with residual endosperm during dissection) and not due to inherent expression of the genes in embryo.

**Supplemental Figure 7.**  Relationships of the temporal regulatory modules to the differentially expressed genes in large- vs. small-sized maize kernels. Differential distribution of the normalized RNA reads (in FPKM) obtained from a developmental series of large (KLS30) vs. small (KSS30) kernels (Sekhon et al., 2014) for each temporal module visualized using a scatter plot of –log_10_-transformed *P*-values. The numbers in each cell indicate the -log_10_*P*-value with a cutoff of *P* < 0.001.

**Supplemental Figure 8.**  Analyses of protein-protein interactions between the candidate TF regulators using yeast two-hybrid assays (Y2H). (A) Yeast *Saccharomyces cerevisiae* AH109 cells were co-transformed with the indicated bait and prey combinations. Proteins listed were fused with Gal4 DNA-binding domain (BD, pGBKT7) or Gal4-activation domain (AD, pGADT7). The combination of empty AD or BD vector with each bait or prey was used as control for self-activation or self-association of bait or prey protein, respectively. Co-transformed colonies growing on QDO were considered to be positive interaction transformants. Plates were incubated at 30°C for 3 days for DDO and QDO. Double drop out (DDO), synthetic complete medium lacking Trp and Leu; Quadruple dropout (QDO), synthetic complete medium lacking Trp, Leu, His, and adenine. (B) A summary table of the Y2H results. The typical E2Fs are highlighted in orange, DPs are highlighted in blue, and atypical E2Fs are highlighted in gray. (C) A graphic representation of the E2F gene network for early endosperm development showing E2F protein interactions and their putative target genes.

**Supplemental Figure 9.** Transactivation and DNA-binding data. (A) The variant/submotifs of E2F-binding sites in M1234 and M123. The number of putative target genes was based on occurrence of the submotifs in -1 kb to +0.5 kb (relative to annotated TSS) of the genes. Because of the possible occurrence of multiple submotifs in the same target gene, the genes corresponding to different submotifs are not mutually exclusive. (B-D) Dual luciferase transactivation assay for the effect of E2F proteins on the different submotifs of E2F-binding site, including the 3× E2F-binding submotif a (B), the 6× E2F-binding submotif b, c, d, and e (C), and the 6× mutated E2F-binding submotif a (TCTCC*aa*C) concatemers (D). Data are fold induction vs. control (mean ± SEM; *n* = 5). Asterisks indicate *P* < 0.05, Student’s *t* test. (E) Electrophoretic mobility shift assays (EMSAs) for binding of E2F16 and E2F6 to upstream sequences of putative target genes containing at least one E2F-binding site (Dataset S11).

**Supplemental Figure 10.** Analysis of transient E2F expression in maize mesophyll protoplasts using RNA-Seq. (A) A summary of the differentially expressed genes (DEGs) resulting from transient E2F expression identified by edgeR (vs. control GFP, FDR < 0.05 and |log_2_FC| > log_2_(1.5)). (B,C) Relative levels of mRNAs visualized using a hierarchically clustered heat map and a boxplot. Asterisks indicate *P* < 0.05, Student’s *t* test. The gene sets that were significantly upregulated in the protoplasts with transient expression of E2F6 in combination with E2F 5 (B) and E2F16 (C) but not E2F6 alone were used for analysis. A mild upregulation of the same genes was detected when E2F5 (B) or E2F16 (C) were singly expressed in protoplasts. (D) Intersection of all ectopic E2F-upregulated genes (E2F-up, see Supplemental Figure 10A), M123-76, and M123 genes without E2F-binding sites in the upstream flanking regions (within -1 kb to +0.5 kb relative to annotated TSS) shown using a Venn diagram. (E) Relative expression of the endogenous *E2F5*, *E2F7* and *E2F13* genes in protoplasts transformed with the E2F16 + E2F6 combo.

**Supplemental Figure 11.**  Analysis of putative Glc-TOR-regulated and KIN10-regulated genes in maize endosperm. (A) The temporal expression patterns of genes encoding TOR complex (TORC), invertases, and sucrose/hexose/sugar transporters during maize endosperm development. Where applicable, the temporal module expression pattern associated with each maize gene is indicated as an extension of the gene name. (B) Overall scheme to determine the maize genes in our dataset that are up/down-regulated by Glc-TOR- dependent processes using the available Arabidopsis data (Baena-Gonzalez et al., 2007; Xiong et al., 2013). As most glucose-regulated genes in WT were shown not to respond to glucose treatment in the *tor* mutant (Xiong et al., 2013), we defined Glc-TOR genes as those responding to the glucose treatment in WT only. (C) Enrichment of Glc-TOR- and KIN10-up/down-regulated genes in our temporal modules visualized using a hierarchically clustered heat map of –log_10_-transformed *P*-values. The numbers in each cell indicate the -log_10_*P*-value and the numbers of genes (in parenthesis) with a *P* < 0.001. *P*-values were calculated using Fisher’s exact test. (D) Enrichment of functional categories of the maize Glc-TOR-regulated genes from the six highest enriched temporal modules for Glc-TOR-upregulated genes (highlighted in yellow) and from the three highest enriched temporal modules for Glc-TOR-downregulated genes (highlighted in green) using MapMan annotation (Thimm et al., 2004), and visualized using a hierarchically clustered heat map of –log_10_-transformed *P*-values (*P* < 0.01). The numbers in each cell indicate the -log_10_*P*-value with a *P* < 0.001 cutoff. *P*-values were calculated by Fisher’s exact test. TCA, tricarboxylic acid cycle. (E) Overall scheme to determine the maize genes in our dataset that are up/down-regulated by KIN10-dependent processes using the available Arabidopsis data ((Baena-Gonzalez et al., 2007; Xiong et al., 2013). (F) Intersection of M123-76 genes, TOR-up genes, and KIN10-down genes shown using a Venn diagram. *P*-values were calculated using Fisher’s exact test.

**SUPPLEMENTAL TABLES**

**Table S1.**  LCM sampling and RNA yields.

**Table S2.**  Reads and mapping statistics of LCM-RNA-Seq data.

**Table** S**3.**  Spearman correlation coefficient analysis of the replicates of the LCM-RNA-Seq libraries.

**Table S4.**  Strategies of preferentially expressed gene identification in endosperm and embryo.

**Table S5.**  *cis*-motifs used for PatMatch analysis of the temporal modules.

**Table S6.** Construction of E2Fs for Yeast two-hybrid assays.

**Table S7.** Construction of E2Fs and *cis*-motifs for dual luciferase report assays.

**Table S8.** Construction of E2Fs for protoplast transient expression assays.

**Table S9.** Construction of E2Fs in E. coli expression for electrophoretic mobility shift assays.

**Table S10.** Sequences of the oligonucleotides used to generate the EMSA probes or competitors.

**SUPPLEMENTAL DATASETS**

**Dataset S1.**  The normalized reads of LCM-RNA-Seq data.

**Dataset S2.**  Dataset of the EN-pref genes.

**Dataset S3**. Dataset of imprinted genes in the maize endosperm.

**Dataset S4.**  Dataset of the EMB-pref genes.

**Dataset S5.**  Dataset of genes obtained from StepMiner analysis.

**Dataset S6.**  Enrichment of TF families in the temporal modules.

**Dataset S7.**  Analysis of *cis*-motifs in the temporal modules.

**Dataset S8.**  Analysis of occurrence of the enriched *cis*-motifs in upstream sequences of the temporal modules.

**Dataset S9.**  Detailed features of the M4-47 network genes.

**Dataset S10.** E2F genes used for phylogenetic analysis in maize, Arabidopsis, and rice.

**Dataset S11.**  Detailed features of the M1234-341 and M123-76 network genes.

**Dataset S12.**  The normalized reads of RNA-Seq data for transient expression in protoplasts.

**Dataset S13.**  The differentially expressed genes of RNA-Seq data for transient expression in protoplasts.

**Dataset S14.**Maize homologs for the Arabidopsis Glc-TOR and KIN10 up/down-regulated genes.

**Dataset S15.** MapMan analysis of maize Glc-TOR-up/down homologs in the temporal modules.

**Dataset S16.** Gene Ontology term enrichment analysis based on Blast2GO and REVIGO.
